# Expanding the Landscape of Disordered Flexible Linkers: A Structural and Computational Framework for DLD dataset assembly

**DOI:** 10.1101/2025.10.26.684646

**Authors:** Di Meng, Juliana Glavina, Heli Magalí García Alvarez, Cesar Osvaldo Leonetti, Gianluca Pollastri, Lucía Beatriz Chemes

## Abstract

Disordered flexible linkers (DFLs) are functional elements found within intrinsically disordered regions that carry out key functions by connecting domains and/or short linear motifs. Understanding the features of DFLs is limited by the lack of comprehensive datasets and accurate predictive models. In this study, we propose a classification for DFLs that includes linkers joining two domains (DLD), a domain and a motif or two short linear motifs. We developed a workflow that allows the systematic identification of DLD-type linkers from protein structures and created a comprehensive dataset known as the DLD dataset. The DLD dataset includes 1640 independent domain linkers (IDLs) which expands currently available linker datasets and annotates related regions such as dependent-domain linkers, intra-domain loops, and termini. Our data collection process integrates missing residue completion and smoothing of short secondary structure stretches enabling to capture a higher number of longer IDLs. We assessed the features of IDLs using t-SNE analysis and protein language model embedding with a CNN-based classifier as well as PCA analysis. IDLs can be distinguished from other disordered and folded protein regions, and their features highly overlap with DisProt Linkers, considered the gold standard for linker annotation. The DLD dataset offers a valuable resource for researchers seeking to investigate the features of disordered flexible linkers and to improve the accuracy and generalizability of DFL predictive models. The DLD dataset is available via an interactive web server at https://dld.chemeslab.org/ where linkers are annotated with sequence and structural features and can be visualized using a structure viewer.

## INTRODUCTION

Intrinsically disordered regions (IDRs) are segments of proteins that lack a stable, well-defined three-dimensional structure under physiological conditions [1,2]. IDRs challenge the structure-function paradigm by playing essential roles in multiple cellular functions despite the lack of a fixed three-dimensional structure [3]. These regions remain dynamic and can adopt multiple conformations, which endows them with functional versatility. IDRs are found in a wide variety of proteins and are particularly abundant in proteins involved in regulation, signalling, and molecular recognition [1–3]. IDRs can be broadly classified based on their functional mechanisms into two categories: binding and non-binding functions [2]. Binding regions within IDRs mediate molecular recognition and signalling by establishing intermolecular interactions with protein partners, whereas a majority of non-binding IDRs mediate entropic functions through their intrinsic disordered states. Among non-binding IDRs, one of the most abundant category is that of disordered flexible linkers (DFLs), which carry out their functions by acting as connectors between functional domains or molecular recognition elements such as short linear motifs facilitating inter-domain communication, multivalent interactions, and contributing to protein structural organization and function [4–9]. According to the latest release of the DisProt database [10] over 75% of non-binding IDRs function as DFLs, highlighting their significance.

Despite their widespread abundance in proteins and the key functional and structural roles they play, the annotation of DFLs remains limited due to the lack of experimental data characterizing disordered linkers, to challenges in distinguishing DFLs from other disordered regions, and to the difficulty in defining their boundaries within diverse protein contexts. Currently, there are only two datasets with annotated DFLs: the DisProt database (DisProt DFL) [10] and the DS-All [11,12] dataset. DisProt [10] is a gold standard database containing IDR annotations that are manually curated from publications that contain experimental evidence for disorder state and functions. DisProt not only provides general IDR annotations but also offers high-level functional classifications related to IDPs, with disordered flexible linkers (DFLs) being a subclass of entropic chains. These entropic chains perform functions directly associated with the disordered state of proteins, such as flexible C-terminal tails, N-terminal tails, and flexible linkers/spacers. DisProt defines flexible linkers as unstructured regions connecting, providing separation, and permitting movement between adjacent functional regions, such as structured domains or disordered motifs [10]. However, despite the growing interest in disordered proteins and their related functions, experimental studies on IDRs remain limited and therefore DisProt is not a comprehensive database, meaning that the majority of IDRs in the human and other proteomes remain unannotated. As of June 2023, DisProt included 338 regions annotated as linkers across 267 protein sequences. Although the DisProt dataset is small, it remains the gold-standard training and evaluation dataset for DFL predictors [13,14] due to its high-quality annotations. The DS-All dataset [11] was originally compiled using domain definitions from the Structural Classification of Proteins (SCOP) [15] database to annotate linkers that connect domains from sequences present in the Protein Data Bank (PDB) [16]. Initially collected with the purpose of improving domain-boundary prediction, DS-All contains 246 linkers from 206 PDB chains [11]. Both DS-All and DisProt contain a similar number of linkers, however DS-All linkers are shorter than DisProt linkers since they are collected from the PDB. Since the length of a linker determines key functional features such as linker flexibility and affinity enhancement in multivalent interactions [9,14], the presence of short linkers is a shortcoming of the DS-All dataset. Several predictors were trained on the DS-All dataset, including two likelihood-based predictors that use amino acid sequence composition and secondary structure prediction [11,12] and three SVM-based predictors [17–19]. However, most of these predictors focused on identifying domain boundaries, and not on comprehensively capturing the linkers between folded domains.

The limited availability of annotated DFLs poses a major challenge for computational studies of linkers, and in particular for developing reliable predictive models. While DisProt remains the gold-standard database for IDRs and DFLs and has been used for training a small number of DFL predictors, its small size introduces bias during model training. Because DisProt annotation depends on the existence of experimental data, annotation of IDRs within protein sequences is not comprehensive. This leads to many linkers present in protein sequences not being annotated, which confounds the definition of negative datasets, reducing the reliability of computational approaches that depend on this data. Small datasets also hinder the identification of key sequence and structural patterns, making it difficult to develop accurate predictors and conduct comparative studies across species and evolutionary contexts. These shortcomings in linker annotation and prediction leave gaps in our understanding of IDR functions, interactions, and dynamics. Although a handful of DFL-specific predictors have shown some improvement through successive Critical Assessment of Intrinsic Disorder (CAID) challenges [20–22], stable and trustworthy predictions of DFLs remain a major challenge. The creation of high quality DFL datasets represents a key step for enhancing predictive models and for gaining a deeper biological understanding of DFLs.

Here, we develop a comprehensive dataset of DFLs tailored for training advanced machine learning models [23]. We outline the methodology for constructing the domain-linker-domain (DLD) dataset. The DLD dataset annotates flexible linkers joining independent domains (IDLs) and joining dependent domains that interact with each other (DDLs). Feature analysis using embeddings and principal component analysis suggest that each of these elements can be separated in a high-dimensional space and have distinct sequence and structural features. We create a dedicated web server interface to explore the features of the DLD dataset using a variety of sequence and structural descriptors as well as the possibility to map linkers to experimental or predicted protein structures. Qualitative and quantitative assessments of the DLD dataset reveals that while these linkers are largely non-overlapping with DisProt linkers from an evolutionary standpoint, they share many features with DisProt Linkers, validating the quality and usability of the DLD dataset for future analysis of linker features and the development of predictive models.

## RESULTS

### Classification of disordered flexible linkers

Intrinsically disordered regions can be distinguished according to their relative location in different structural and/or functional elements of a protein sequence. In this work, we introduce a three-class classification scheme for disordered flexible linkers or DFLs (Figure 1A). DFLs in the Domain-Linker-Domain (DLD) class can connect two adjacent but independent folded domains (independent domain linker, IDL) or two adjacent and structurally dependent domains that establish intramolecular interactions (Dependent Domain Linker, DDL). DFLs can also belong to the Motif-Linker-Domain (MLD) class, where the DFL connects a short linear motif to a globular folded domain, facilitating domain-motif interactions. Finally, DFLs can belong to the Motif-Linker-Motif (MLM) class, where the DFL connects two short linear motifs, enabling flexibility and multivalency in motif-based interactions. DFLs differ from flexible loops (L) that connect structural elements within a folded domain and from disordered Terminal (T) regions located at the N- or C-terminus of a protein (Figure 1B). For the remainder of this work, we focus on linkers connecting folded domains (DLD class).

**Figure 1.**
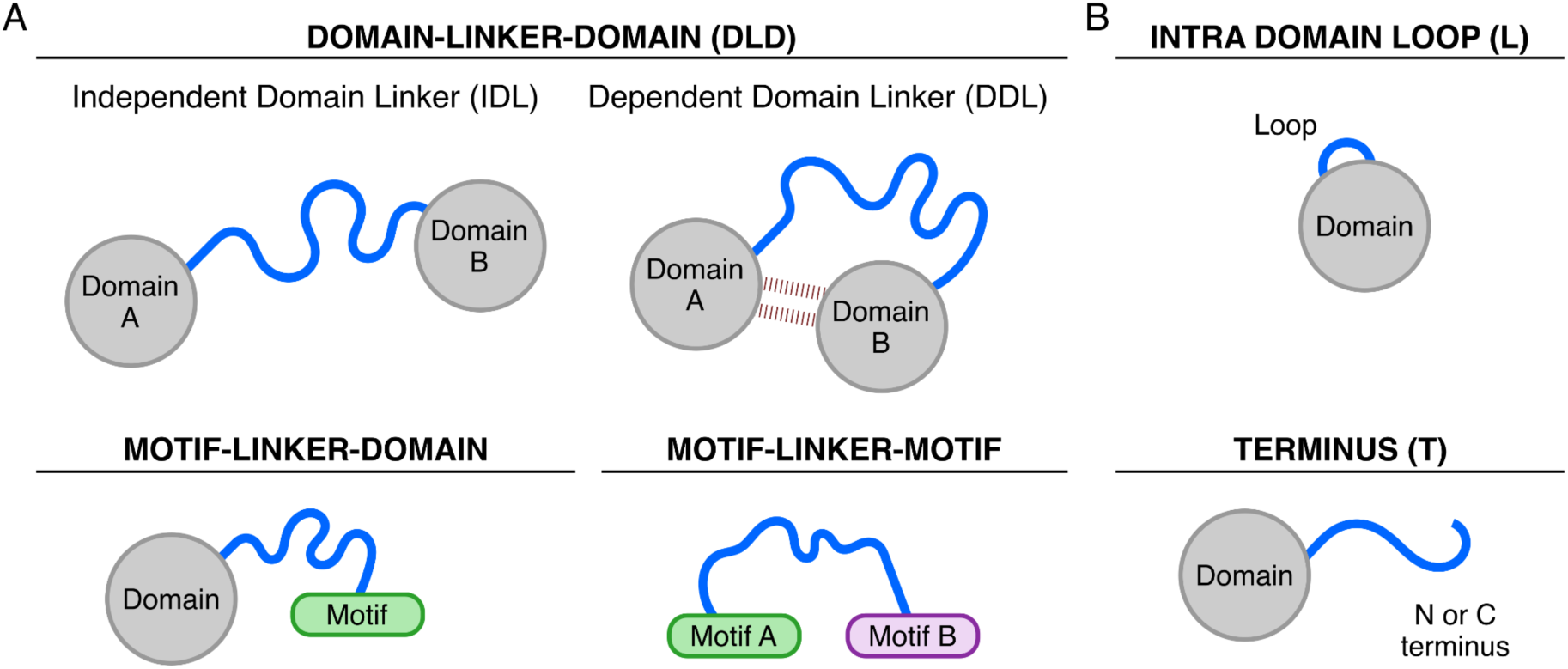
Classification of Disordered Flexible Linkers (DFLs), Loops and Termini. (A) Classes of DFLs. The Domain Linker Domain (DLD) class includes Independent Domain Linkers (IDL) and Dependent Domain Linkers (DDL). Motif Linker Domain and Domain Linker Motif topologies are grouped in the MLD class. The third is the Motif Linker Motif class. (B) Loops are defined here as flexible regions connecting secondary structure elements within a folded domain. Termini are defined as intrinsically disordered regions at the N- or C-terminus of a protein. For the DLD dataset, termini represent a flexible region at the N- or C-terminus of a domain and in most cases do not represent the true N- or C-terminus of the protein. Domains are depicted as grey circles, motifs are depicted as colored rounded rectangles, and DFLs and Loops are depicted as blue lines.

### DLD dataset construction

To construct a high-quality dataset of DFLs connecting folded domains (DLD-class), we developed a systematic pipeline to identify these linkers and classify them as IDLs or DDLs (Figure 1A). The pipeline was inspired by the DS-All dataset [11] but involved multiple improvements. First, the size of the starting dataset was increased from 206 PDB chains containing multidomain proteins in the DS-All dataset to 2712 PDBs in the DLD dataset. Second, our method improved the detection of linker boundaries through missing residue completion and two smoothing steps. A detailed overview of the data collection process is presented in Figure 2, Supplementary Figure S1 and Materials and Methods.

**Figure 2.**
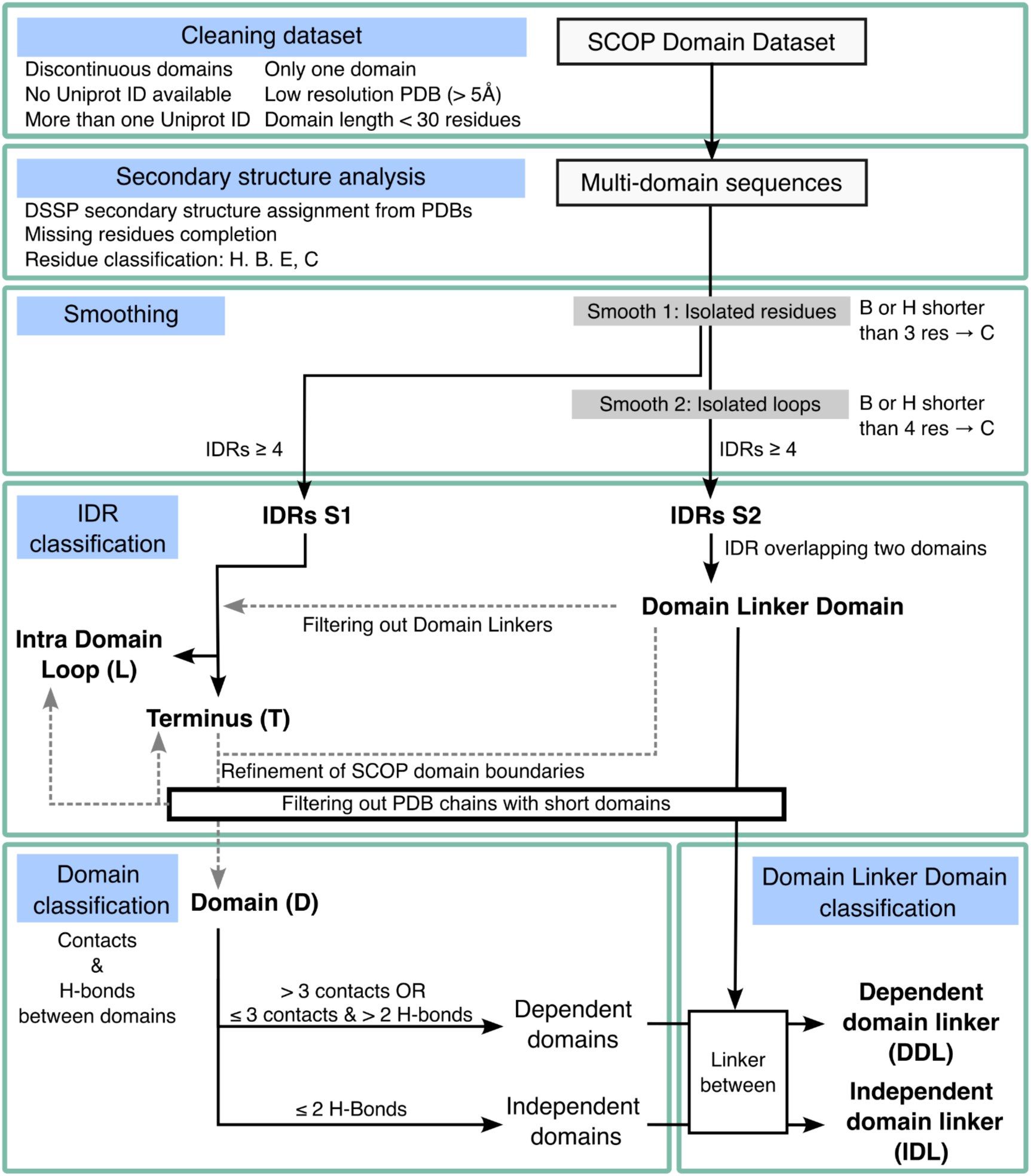
Summary of the construction process of the DLD dataset. See Supplementary Figure S1 for an expanded version.

We selected the SCOP database for domain annotation due to its structural emphasis, manual curation, and comprehensive, residue-level coverage [15,24]. SCOP provides high-quality annotations of protein domains based on a structure-centric classification of experimentally determined protein structures obtained from the Protein Data Bank (PDB) [25]. First, using ∼30000 PDBs from the SCOP2 dataset as a starting point, we performed a filtering step to retain PDBs containing multidomain constructs and discarded PDBs containing domains with non-continuous topologies, chimeras, low quality structures or very small (<30 res) domains, which led to the annotation of SCOP domain boundaries across 2712 PDBs containing at least two protein domains (Figure 2).

Second, we annotated a structural category (helix, beta or coil) to each residue using DSSP [26,27] and a reclassification of the eight DSSP states into four states (see Materials and Methods). To better capture the full extent of domain linkers, we applied two preprocessing steps. First, we completed missing residues from the PDB chain which are often disordered in crystal structures and annotated them within DSSP group C. Second, we performed a two-step smoothing process to remove short (< three or four residue) stretches of local secondary structure present within linker regions. This allowed extending IDRs while preserving the structural integrity of folded domains. Following all preprocessing steps, we performed a reclassification of domain boundaries.

Finally, we annotated IDRs connecting two adjacent domains as domain linkers. These linkers were classified into two categories based on the degree of structural interaction between the domains. Independent domains shared minimal inter-domain contacts (≤3 C–C contacts and ≤2 hydrogen bonds), and linkers between them were labelled as IDLs. Linkers connecting domains with a higher number of interactions were categorised as DDLs.

The DLD dataset contains 1640 IDLs and 647 DDLs (Table 1). In addition, the pipeline led to the annotation of a high number of intra-domain loops (L), termini (T), and domains (D) (Table 1), resulting in a structurally grounded dataset.

**Table 1.**
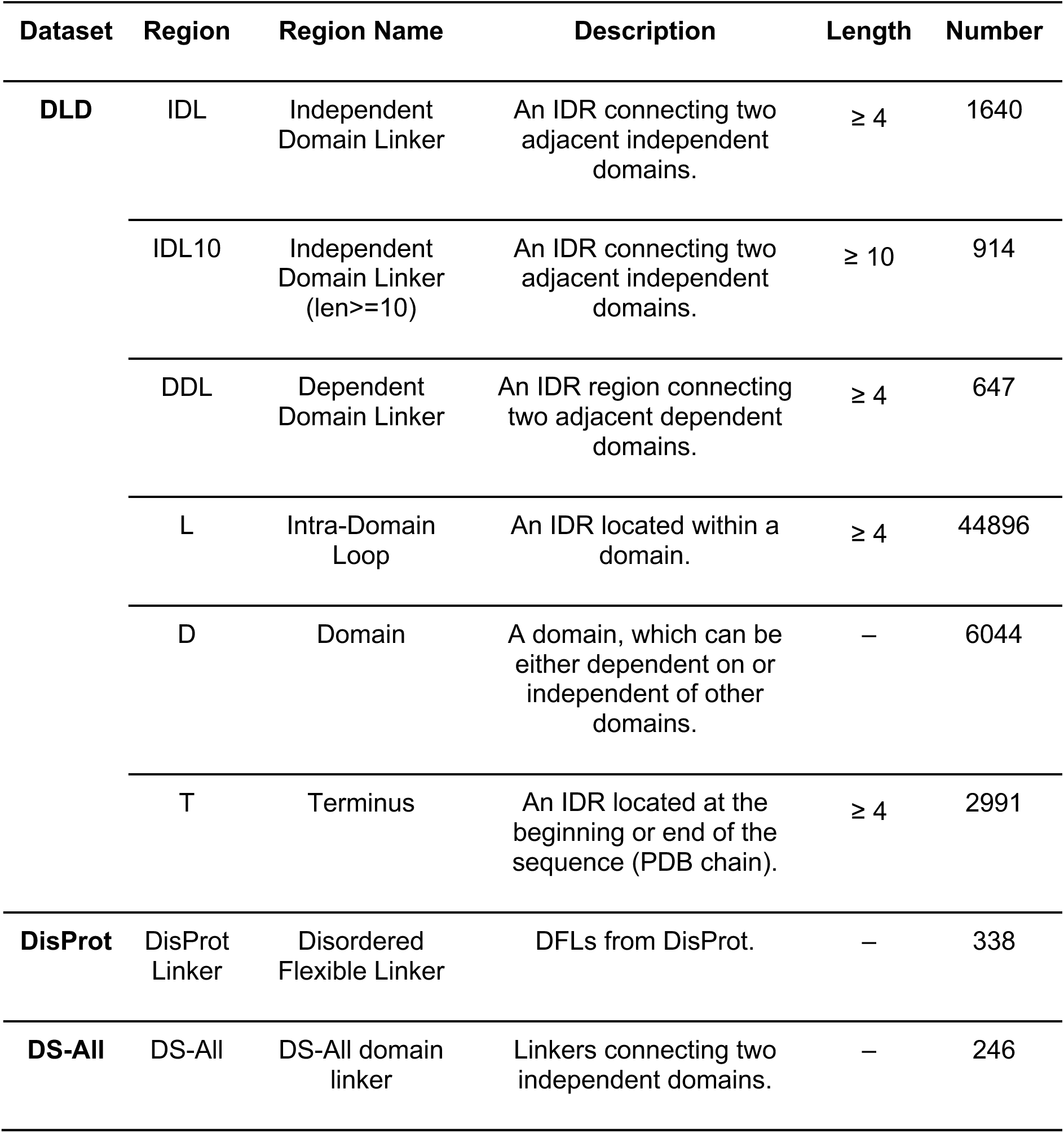
Summary of the datasets and relevant information and description of regions.

### Improved Coverage and Linker Length

The DS-All dataset [11] was developed with the primary aim of improving domain boundary prediction [12,17–19]. In contrast, our pipeline focused on comprehensively capturing domain linkers. One limitation in annotating longer linkers from the experimental structures is the presence of missing residues and/or short stretches of structured residues. Two key enhancements were critical to overcome this: (1) missing residue completion in protein structures, and (2) refinement of structural annotations by a two-step smoothing process. Together with the use of a much larger dataset of PDBs as a starting point when compared to previous datasets, these improvements enabled the identification of more numerous and longer IDLs.

A representative example of the identification of linkers through the annotation of missing residues is shown for the Seven less Son 1 (Supplementary Figure S2A). When applied across all PDBs, this procedure did not lead to annotation of new IDLs but did increase the length for ∼10% of IDLs, since 152 of the 1640 IDLs annotated in the final dataset contain missing residues (Supplementary Figure S2B). The effect of missing residue completion was most notable for longer linkers, representing roughly 23% of annotations for linkers ranging from 20-30 residue length, 30% of annotations for linkers ranging from 30-40 residue length and 50% for linkers ranging from 40-70 residue length (Supplementary Figure S2B). This demonstrates that filling in the gaps helps to extend IDLs beyond what is typically captured in raw structural data, especially improving the representation of longer IDLs.

The two-step smoothing process further enhanced annotation quality by removing short, isolated stretches of secondary structure that otherwise fragment linkers leading to missing these annotations (Figure 3). Figure 3A exemplifies the effect of the two smoothing steps (see Materials and Methods). For the 2BNM_a chain, the first step of smoothing extends the length of the linker from 8 to 16 residues, while the second step of smoothing extends the length of the linker from 16 to 29 residues. For the 4B2Y_a chain Figure 3B, only the second step of smoothing significantly extends the length of the linker, from 9 to 34 residues, indicating that it is possible that the IDL length remains unchanged after one smoothing step and that the linker was already properly captured during the initial smoothing. The two-step smoothing process led to capturing an additional 210 IDLs (Figure 3C, Table 2) and extended the length of 270 linkers. As a result, smoothing increased the number of linkers of length 10 and higher by 307 (Figure 3C). Both smoothing steps are necessary to capture more and longer linkers (Table 2). When considering all linker classes, the smoothing led to annotating 272 additional linkers, improving the annotation of IDLs and DDLs from 2015 to 2287 (Table 2).

**Figure 3.**
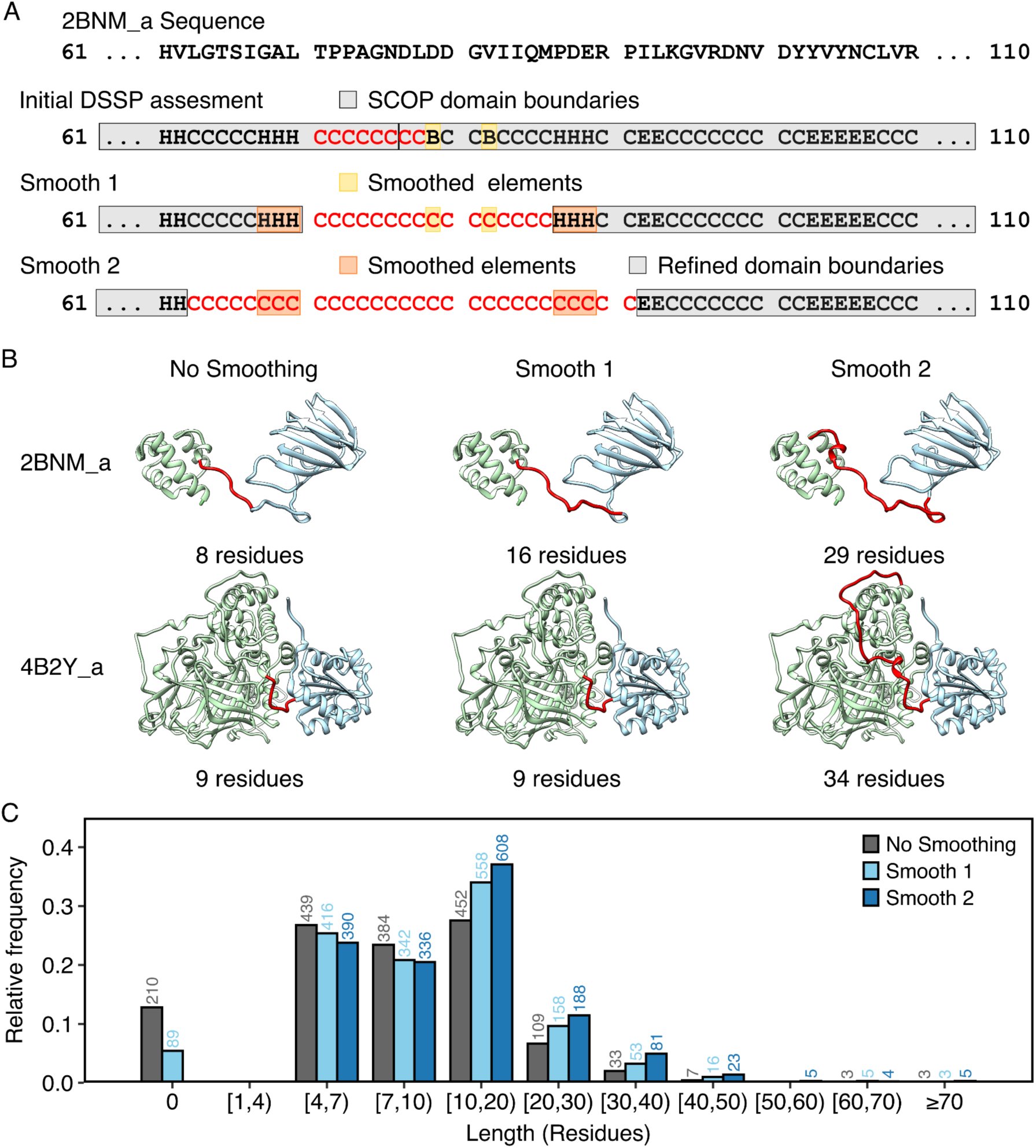
Effect of the number of smoothing steps on Independent Domain Linker length. (A) Sequence and secondary structure assignment by DSSP and after one or two smoothing steps on a fragment of the PDB 2BNM_a. Domain boundaries are indicated in gray, starting from SCOP domain boundaries to the final refined domain boundaries. Darker colours highlight isolated residues assigned as helices or beta sheets that are going to be smoothed. The detected IDL is highlighted in red. (B) The PDB chains are oriented from the N-terminal (left) to the C-terminal (right). Domains are colored in gray (if present), green and blue from the N-terminal to the C-terminal. IDLs are highlighted in red and orange if present. Dashed lines indicate missing residues. (C) Linker length distribution with No Smoothing and one or two smoothing steps (Smooth 1 or Smooth 2 respectively). A linker length of 0 residues long means that these linkers were not detected on the corresponding smoothing step.

**Table 2.**
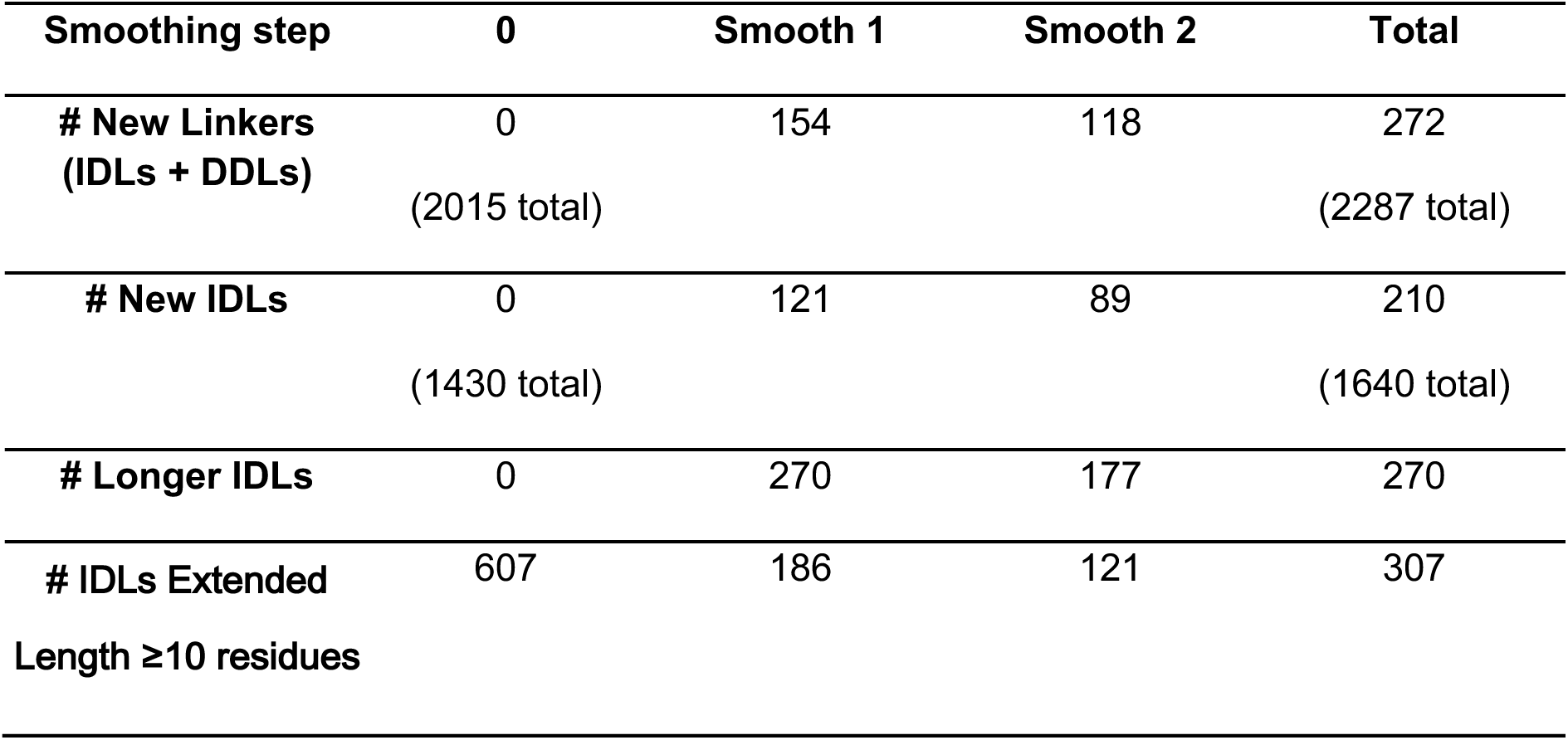
Effect of two-step smoothing on the number of domain linkers.

Overall, these results highlight the effectiveness of the two-step smoothing process and the completion of missing residues in producing a more accurate representation of IDLs and improving the identification of longer linkers.

### Comparison of the DLD dataset to other linker datasets

To assess the DLD dataset, we compared the features of IDLs and DDLs to linkers from the DS-All dataset [11], which was generated from PDB structures using computational analysis and to IDRs annotated as Linkers (class IDPO:00502) in DisProt, a reference dataset considered the gold standard in the field where IDRs are curated from experimental evidence and linkers are defined as a functional class following an IDR-specific Ontology [11]. The total number of IDLs in the DLD dataset (1640) improves the number of linkers annotated in DS-All and DisProt by 7-fold and 5-fold respectively (Table 1 and Figure 4A). When compared to DS-All and DisProt (Figure 4A and Supplementary Figure S3), the DLD dataset contains more linkers across most length bins, with DisProt only surpassing DLD for long length bins (linkers ≥ 40 residues, Figure 4A). DS-All identifies only 34 linkers longer than 20 residues, while the DLD dataset identifies 306 linkers in the same length category. These results highlight that our pipeline improved the annotation of long IDLs even when compared to reference databases such as DisProt, which place high efforts on annotating long IDRs.

**Figure 4.**
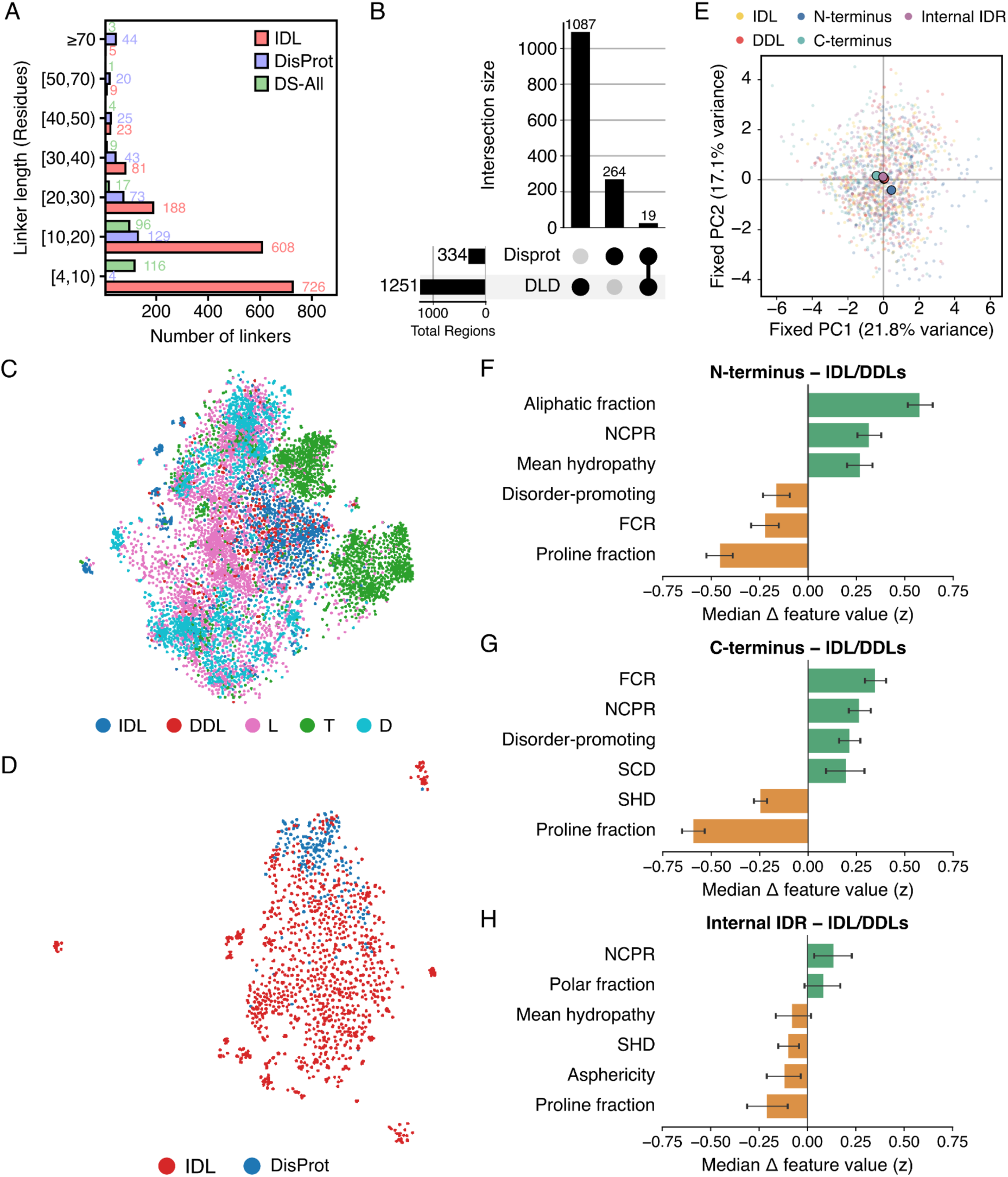
Characterisation of Linkers, spatial visualisation and comparison of IDRs and linkers. (A) Length distributions of the three linker datasets: IDL, DS-ALL, and DisProt Linker. (B) Upset plot showing the number of DLD-specific, DisProt-specific and overlapping linker region clusters. The datasets were filtered for regions > 9 residues and the thresholds used were 20% identity and 30% coverage. The side bar graph indicates the number of regions for each dataset. (C) t-SNE plot of the 5 regions in the DLD dataset. See Figure Supplementary 2 for pairwise comparison t-SNE plots. (D) Pair comparison between the IDL and DisProt Linker datasets. See Figure Supplementary 3 for all DLD regions - DisProt comparison t-SNE plots. DLD: Domain-Linker-Domain. IDL: Independent domain linker. DDL: Dependent domain linker. L: Intra domain loops. T: Terminus. D: Domains. (E) PC1–PC2 projection of a representative resample onto axes defined by PCA of pooled length-matched regions across all resamples. Included region classes were IDLs, DDLs, N-termini, C-termini, and Internal IDRs. Length matching yielded n = 288 regions per class (n = 1,440 per resample). Points represent individual regions and outlined circles indicate class centroids. The displayed resample was selected as the one whose ten pairwise PC1–PC2 centroid distances among the five plotted classes were closest to the median distance profile across 500 resamples. (F–G) Median differences (*Δ*) in standardized feature values (z) for N-termini, C-termini, or Internal IDRs relative to the equally weighted pooled IDL/DDLs group across 500 length-matched resamples. Positive values indicate higher feature values in the indicated class than in structural linkers. Error bars show the empirical 2.5th–97.5th percentile range across resamples. Each panel shows the six features with the largest absolute median difference for the corresponding comparison.

To provide a stringent assessment of sequence similarity between the DLD and DisProt datasets for linkers longer than 9 residues, we performed clustering analysis at 20% sequence identity and 30% sequence coverage (Materials and Methods). At the protein level, only 9% of the protein clusters containing DLD sequences also contained DisProt sequences (Supplementary Table S1 and Figure S4). To provide a more accurate assessment of the overlap between linker annotations, we conducted the same analysis at the region level. At the region level, only 1.7% of the clusters (19 clusters) containing DLD sequences were shared with DisProt (Figure 4B and Supplementary Table S1). Therefore, using stringent clustering criteria reveals that the DLD dataset is largely non-overlapping with the DisProt dataset both at the protein and at the linker region levels. Inspecting DisProt and DLD annotations for the 19 shared clusters (Supplementary Figure S5) revealed that, in 15 out of 19 cases DLD identifies regions that are longer or equal to those annotated based on experimental evidence in DisProt. Within the limits of this small dataset, these results suggest that the DLD pipeline can identify slightly longer regions, achieving a more complete coverage of the end points of linker regions compared to DisProt.

The DLD dataset contains ∼1000 short IDLs/DDLs (<10 residues, Supplementary Figure S3). One possibility is that the pipeline annotates short linker regions corresponding to linkers connecting the structural units of repeat domains. To assess this possibility, we inspected the number of short linkers per PDB chain. The large majority (87%) of PDB chains contain only one short linker (Supplementary Figure S6), suggesting the short linkers do not map between repeat units. Moreover, only 57 of 951 proteins containing a short IDL/DDL (6%) contain a repeat domain annotated by Pfam, and only six linkers map within or between these repeat domains (Supplementary Figure S7). This result indicates that the abundance of short linkers in the DLD dataset is not driven by a systematic inclusion of linkers connecting the individual units of repeat domains.

### Features of the DLD Dataset

To perform a first assessment of the features of DLD linkers, we compared their amino acid composition to that of DisProt and DS-All Linkers (Supplementary Figure S8). All datasets showed a sequence composition characteristic of disordered regions with an enrichment in polar and charged amino acids and in structure-breaking amino acids such as glycine and proline, with slight differences such as DisProt being more enriched in serine and glutamine and IDLs being more enriched in proline (Supplementary Figure S8). DS-All linkers showed nearly identical profiles to IDLs and DDLs from the DLD dataset (Supplementary Figure S8).

Next, we employed ProtTrans [28] to embed DLD dataset sequences corresponding to different structural classes in a high-dimensional feature space and used t-SNE (t-Distributed Stochastic Neighbour Embedding) [29] to project the high dimensional sequence embedding data into two dimensions. The t-SNE analysis revealed that IDLs clustered closely with DDLs but remained distinct from domains (D), loops (L), and termini (T) (Figure 4C). Pairwise comparisons reinforced this finding (Supplementary Figure S9). Some loops overlapped with IDLs while most did not, which is likely to reflect differences between a smaller proportion of longer loops that are more similar to linkers in properties and a larger proportion of short loops which are more structured (Supplementary Figure S9). Termini clustered largely apart from IDLs, indicating the presence of distinct properties for this class.

The comparison of IDLs and DDLs to DisProt linkers showed a higher overlap between IDLs and DisProt linkers (Figure 4D) and a comparatively lower overlap between DDLs and DisProt linkers (Supplementary Figure S10). This result is of special interest since it emphasizes that the DLD pipeline captures linkers with features that align with those found in gold standard datasets such as DisProt. However, IDLs are distinct from DisProt linkers from an evolutionary standpoint (Figure 4B, Supplementary Table S1 and Figures S4 and S5), indicating that the DLD pipeline expands available datasets with high-quality linker sequences.

To identify interpretable sequence- and structure-derived properties associated with the similarities and differences among region types, we performed a principal component analysis using 13 non-redundant features (see Materials and Methods, Supplementary Text, and Supplementary Figure S11). Across all region types, domains occupied the hydropathy-, aliphatic fraction-, and sequence hydropathy decoration (SHD)-associated end of PC1, whereas the non-domain classes were shifted toward the opposite end and substantially overlapped (Supplementary Figure S11B). Thus, these features captured the broad distinction between ordered and disordered regions but did not clearly resolve the different linker subclasses. When comparing length-matched IDLs, DDLs, and DisProt linkers (Supplementary Figure S11C), the PC1 axis separating DisProt linkers from IDLs and DDLs was defined primarily by the fraction of charged residues and disorder-promoting composition at one end, and SHD, mean hydropathy, scaled radius of gyration, and aliphatic fraction at the other. This pattern is consistent with the compositional analysis (Supplementary Figure S8), which showed that DisProt linkers were relatively enriched in serine and glutamine, both of which contribute to the disorder-promoting amino acid fraction and suggests some differences in compaction between DisProt linkers and IDLs/DDLs.

The broad termini class comprises both bona fide protein N- and C-termini and internally located regions arising from crystallographic construct boundaries. Since this heterogeneous class clustered largely apart from other regions in the t-SNE analysis (Figure 4C), we next asked whether this pattern reflected features specific to true termini, construct-derived Internal IDRs, or both. We therefore analyzed N-termini, C-termini, and Internal IDRs separately, comparing each category with an equally weighted IDL/DDLs reference group across length-matched resamples (refer to Supplementary Text for methods).

In the fixed PC1–PC2 projection, Internal IDRs overlapped substantially with IDLs and DDLs and had a centroid close to the IDL/DDL groups (Figure 4E). By contrast, N- and C-termini were displaced from DL/DDLs and showed modest separation from one another. However, the broad overlap among individual regions suggests shifts in average feature profiles rather than completely distinct populations.

To identify the features underlying these shifts, we calculated the standardized difference for every feature between each category and the IDL/DDLs reference group in every resample. Figure 4F–H shows the six features with the largest absolute median differences for each comparison. Relative to IDL/DDLs, N-termini showed higher aliphatic fraction and lower proline fraction, together with moderately higher NCPR (Figure 4F). The increased aliphatic fraction is consistent with a contribution from the initiating methionine at protein N-termini, as also suggested by Supplementary Figure S8B. C-termini were primarily distinguished by lower proline fraction and higher FCR and NCPR (Figure 4G). This was in agreement with the higher relative frequency of lysine and glutamic acid in C-termini compared to IDLs (Supplementary Figure S8C). Internal IDRs showed smaller contrasts overall (Figure 4H).

While the PC1–PC2 projection captured the relative positioning of the classes, it represented only part of the feature contrasts (Figure S12A). The first two PCs captured median fractions of 43.6%, 29.0%, and 9.1% for N-terminal, C-terminal, and Internal IDR contrasts, respectively; after inclusion of PC5, cumulative capture reached 90.1%, 83.2%, and 80.1%. The contribution heatmaps showed that the three comparisons were represented by different combinations of fixed PCs rather than by a single common terminal-versus-linker axis (Figure S12B–E). Thus, the differences between termini and IDLs/DDLs are multidimensional and only partly revealed by the PC1–PC2 projection.

In sum, the sequence composition and PCA analyses provide interpretable clues about the sequence and structural features that distinguish region types but they do not fully explain the similarities and differences captured by the ProtTrans embeddings. This highlights that relating embedding-derived separation to specific biological or biophysical determinants remains an open challenge that will require further investigation.

### Quantitative classification of similarity between different regions identified by the DLD dataset and reference linker databases

While t-SNE plots provide an intuitive 2D visualization of non-linear relationships between regions independent of region length, they lack a quantitative measure of similarity between regions. To address this limitation, we complemented t-SNE analysis with a classifier-based analysis. For this, we trained classifiers based on a simple M2O neural network to distinguish between region types (Supplementary Figure 13). The performance of these predictors is inversely correlated to the similarity between regions

Our assessment focused on comparing the performance of the predictors in two groups. The IDL group (Group 1) trained the predictor to distinguish between IDLs and the rest of the DLD dataset regions (DDL, L, D and T), whereas the DisProt linker group (Group 2) trained the predictor to distinguish between DisProt linkers and all DLD dataset regions (IDL, DDL, L, D and T). We separated Group 1 and Group 2 comparisons because different DLD regions may originate from the same sequence, introducing a level of similarity that can make them harder to distinguish. This systematic bias affects all predictors in Group 1 similarly, allowing us to ignore it in our comparisons. However, since DisProt Linker is not subject to this bias, we cannot directly compare predictors that classify DisProt Linker against IDL with those that classify IDL against DDL. Instead, we compare predictors that classify DisProt Linker against all DLD regions separately.

All predictors were trained for the number of epochs required to reach convergence, and the results are summarized in Supplementary Table S2. In Group 1, the lowest performance (AUC score 0.762) was obtained for the predictor that distinguishes IDLs from DDLs (pIDL_DDL), whereas the other three predictors achieved AUC scores above 0.96. This indicates that IDLs are more challenging to distinguish from DDL than from loops (L), domains (D) or termini (T), confirming the t-SNE and PCA findings that IDLs and DDLs are the most similar regions. The highest AUC score in Group 1 is from the predictor that separates IDL from D (pIDL_D), as expected from the clearly different sequence composition of folded domains (D) and IDR regions (Supplementary Figure S8) [30].

When comparing DisProt Linker against all DLD regions, pIDPO_IDL achieved a lower AUC score (0.875) compared to the other four predictors in Group 2, indicating that DisProt Linkers and IDL are more similar to each other than DisProt Linkers are to any other DLD regions. DDL were distinguished from DisProt linkers with higher performance than IDL, indicating in agreement with t-SNE plots that IDLs are the most similar region to DisProt linkers. This may be due to the higher flexibility of linkers joining non-interacting domains as compared to interacting domains with a fixed spatial orientation to each other.

Altogether, t-SNE and quantitative classifier analysis confirm that IDLs present the strongest sequence and feature similarity with DisProt linkers followed by DDLs, and that IDLs/DDLs remain distinguishable from other protein regions.

### DLD-Dataset Web server

To facilitate the exploration of the DLD Dataset we develop a dedicated web server with an intuitive user interface (https://dld.chemeslab.org/). The DLD dataset web server allows the user to explore the IDLs, DDLs, Termini and DisProt linkers through an interface organized into two main tabs, “Main” and “Viewer” (Figure 5).

**Figure 5.**
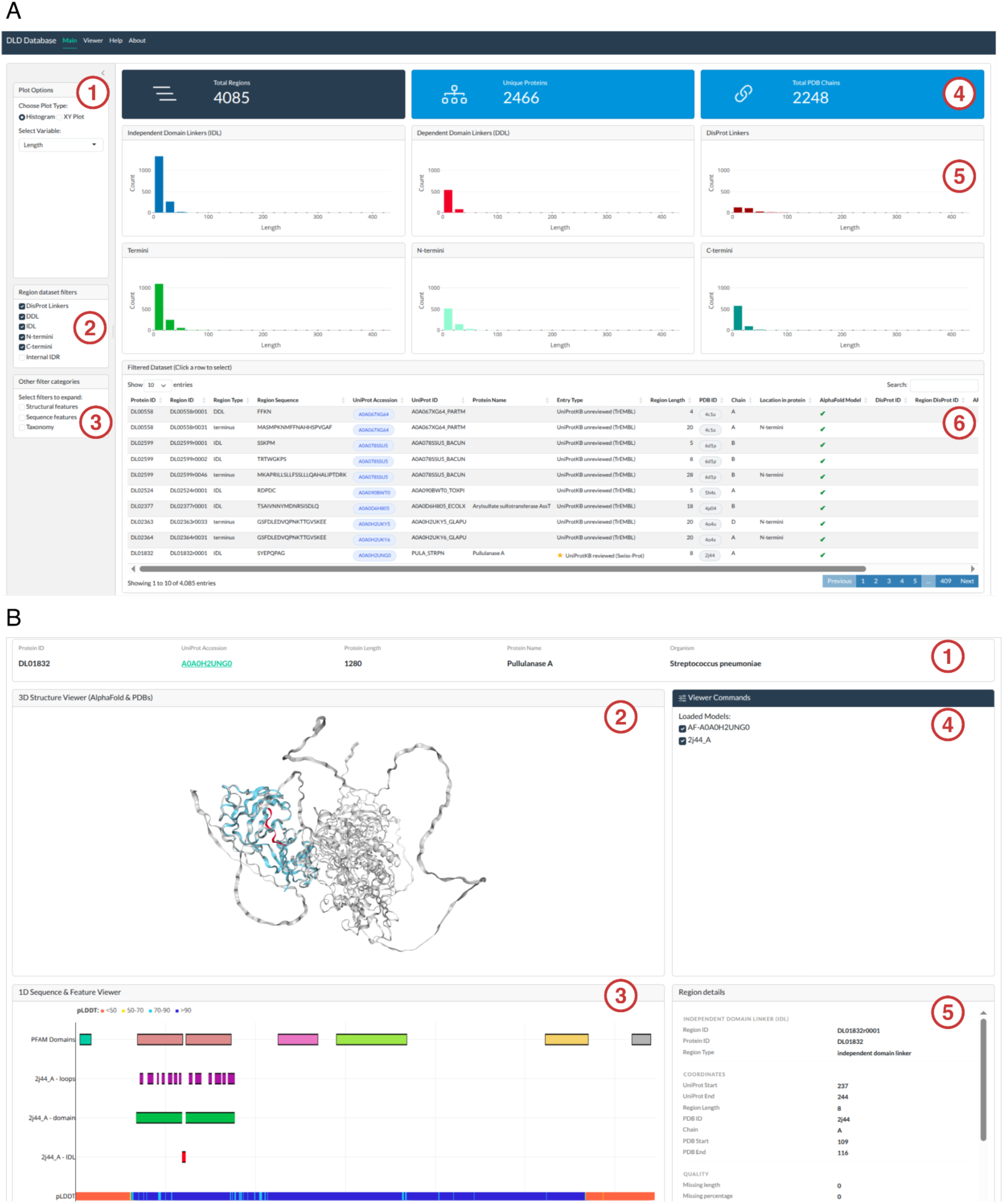
DLD-Dataset Web Server. (A) The Main tab provides an overview of the database and interactive exploration tools. The left side bar includes three panels: (1) plot options to toggle between histogram and XY plot views and select the feature/s displayed, (2) region dataset filters allow users to filter by region type (DisProt linkers, DDL, IDL) and (3) additional expandable filter categories for structural features, sequence features, and taxonomy. On the right, (4) summary cards display the total number of regions, unique proteins and PDB chains in the filtered dataset, (5) interactive graphs show the distributions or scatterplots of the selected feature/s and (6) the data table presents the filtered dataset; rows can be clicked to load a protein into the Viewer. (B) The Viewer tab allows inspection of individual proteins and their linker regions. (1) The top bar displays the protein annotations, (2) an interactive 3D structure viewer shows the AlphaFold model and the PDB chain superimposed when available, with visibility controlled via the Viewer Commands panel (3). The sequence tracker (4) displays Pfam domain annotations, DisProt linker evidence, DLD loops, domains and IDL/DDL linkers, and the pLDDT scores. Clicking on a region highlights it in the structure and loads additional information in the region details panel (5).

The “Main tab” (Figure 5A) is organized in two areas. The left side bar contains three panels with plot controls (1) and interactive filters (2 y 3). The plot options panel (1) allows the user to select between distribution and XY plot views and to select the feature or features to plot. The region dataset filters panel (2) allows the user to restrict the data by region type: IDLs, DDLs, N-termini, C-termini, or DisProt linkers. An additional set of expandable filters categories (3) enables filtering by sequence features like (e.g. NCPR, FCR), structural features (e.g. predicted end-to-end distance,radius of gyration) and taxonomy groups. On the right side, three panels update dynamically in response to the applied filters. The summary cards (4) display the total number of regions, unique proteins and PDB chains in the filter datasets. The interactive graphs panel (5) displays the distribution of a selected feature or the scatterplot of two selected features for each region type. Last, the bottom panel (6) includes a filterable and searchable data table that presents the filtered dataset. Each row of the table corresponds to a region and can be clicked to load the corresponding protein into the Viewer.

The “Viewer tab” (Figure 5B) allows the inspection of individual proteins and their associated linker regions. The top bar (1) displays protein annotations including, the accession number and protein name. The central panel provides an interactive 3D structure viewer (2) that displays when available the AlphaFold model and the PDB chain used in the construction of the DLD dataset. The individual models can be toggled on/off via the Viewer Command panel (3). At the bottom, an interactive sequence tracker (4) displays when available Pfam domain annotations, DisProt linker evidences, DLD Dataset associated regions (loops, domains, IDLs, DDLs, termini) and the pLDDT scores obtained from the AlphaFold model. Clicking on any region highlights the region in the 3D structure viewer and the panel “Sequence Details” (5) will show the associated features to that region. Finally, the raw data used to construct the DLD dataset and processed data underlying the web server are available for download from the “Help” tab.

## DISCUSSION

The annotation and characterization of IDRs that function as linkers will improve our understanding of the features of this functional class of IDRs. While some efforts have been made to develop linker predictors using available data [11–14,17–22], the generation of comprehensive datasets of DFLs is a necessary step for the refinement of prediction tools for disordered linkers.

Disordered flexible linkers (DFLs) can be broadly classified into linkers joining folded domains (DLD), a domain and a short linear motif (MLD/DLM) and linkers joining two short linear motifs (MLM). Folded domains can be classified based on structural analysis [24,31–34] and multiple computational tools are available to identify domain families [35–38], making domains easier to identify compared to short linear motifs [39–44]. Building on this advantage, we developed a workflow that identifies DLD-type linkers from protein structures. The DLD dataset includes 1640 independent domain linkers (IDLs), 647 dependent domain linkers (DDLs) and other related protein regions, such as intra-domain loops (L), and termini (T). Our data collection process integrates missing residue completion and smoothing of short stretches within linkers which improves the extraction of longer IDLs. The methodology is scalable for its application to larger datasets of protein structures.

Our analysis of the DLD dataset revealed that IDLs possess distinct characteristics compared to other protein regions, including intra-domain loops, domains, and termini. By applying machine learning methods, including sequence embeddings from ProtTrans, convolutional neural networks, and visualization with t-SNE, we identified structural and sequence similarities between IDLs in the DLD dataset and DisProt linkers. Clustering analyses confirmed that while the DLD pipeline extracts linkers with similar sequence features to those of gold standard DisProt linkers, while representing distinct sequences from an evolutionary standpoint. These results demonstrate the robustness of our data collection strategy and its usefulness for expanding current linker datasets.

The DLD dataset offers a valuable resource for researchers seeking to improve DFL prediction accuracy and generalizability and to develop computational methods to assess DFL features. Looking ahead, the DLD dataset can be expanded by integrating a larger number of structures from the PDB and predicted structures from the AlphaFold database [45,46]. This will enable the annotation of a much larger dataset of IDLs. The development of methods that allow identifying other linker types such as DLM/MLD and MLM, will further advance our understanding of DFLs and their roles in protein function. This work contributes towards the continued development of DFL-specific prediction tools and provides a resource for researchers aiming to explore the structural and functional roles of disordered flexible linkers in proteins. The DLD dataset is made available through a dedicated web server (https://dld.chemeslab.org/) that allows users to interactively explore, filter, and visualize IDLs, DDLs, termini, and DisProt linkers alongside an array of sequence and structure descriptors.

## MATERIALS AND METHODS

### Domain Dataset collection and filtering

To create a dataset of domain linkers we downloaded all structurally defined domain annotations from SCOP2 released on June 29, 2022 [24] as a text file (Supplementary File S1). We adopted the family-level classification from SCOP2, which is based on structural similarity as the domain definition. We collected annotations for 31477 PDB chains containing 36003 domains from 30324 UniProt sequences.

Several filtering criteria were applied for cleaning the initial dataset. First, to ensure continuity between adjacent domains [47], we excluded 382 PDB chains containing discontinuous domain topologies that mapped to multiple non-contiguous segments of a UniProt sequence (FA UNIREG column in Supplementary File S1). Second, we excluded 93 PDB chains that could not be mapped to a single UniProt accession number and possibly represented chimeric proteins. Last, we filtered out 28135 single-domain PDB chains to create a dataset of PDB chains containing multiple domains. As a result, we retained 2867 multi-domain PDB chains for further analysis. We downloaded 2734 mmCif files representing the 2867 multi-domain PDB chains from PDBe [48] on March 11, 2023. To improve the quality of our dataset we filtered out 155 PDB chains that had low resolution (resolution > 5 Å) or that had at least one domain shorter than 30 residues obtaining a final dataset of 2712 PDB chains. A detailed overview of the SCOP2 data processing steps is provided in Figure 2 and expanded in Supplementary Figure S1. The final dataset was compiled as a table containing essential information such as PDB ID, chain ID, domain region start and end positions, and domain family ID (Supplementary File S2).

### Secondary structure annotation

To parse the data contained within the PDB chains into folded and disordered regions, we used secondary structure annotation. Specifically, we used DSSP v4.0.4 implemented in BioPhyton v1.80 to assign secondary structure at the residue level on the 2712 PDB chains. DSSP could not parse 47 mmCif files representing 63 PDB chains (Supplementary Figure S1), resulting in 2649 PDB chains containing DSSP assignments. We adopted a 4-state classification system that grouped the eight states defined by DSSP. The classification scheme we used consisted of states H (helix), B (isolated beta strand), E (extended beta strand) and C (Coil). The state H included the DSSP states H (Alpha helix), G (3-10 helix) and I (π-helix). The state B included the DSSP state B (Isolated beta bridge). State E included the DSSP state E (beta Strand) and state C included the DSSP states T (Turn), S (Bend) and C (residues where no structural state was assigned by DSSP). The DSSP algorithm does not assign a secondary structure class to missing residues in PDB files, which often correspond to intrinsically disordered regions including loops and linkers. In order to improve the annotation of disordered regions in the PDB files, we extracted information on missing residues from mmCif files and labeled them as state C.

### Identification of disordered regions

The identification and classification of intrinsically disordered regions within PDB files required adopting a definition based on the DSSP and SCOP2 domain annotations. We defined an intrinsically disordered region (IDR) as a region containing at least four contiguous residues classified as Coil (C) that lack regular secondary structure, defined by categories H, B and E. To create a functional classification of IDRs, we further distinguished between IDRs linking adjacent domains (interdomain linkers) and IDRs located within a domain (intradomain loops and terminal regions or termini) (Figure 2). Domain linkers were defined as IDRs that overlapped with two SCOP2 domain boundaries and included continuous residues assigned a C state, including those overlapping with the SCOP2 domain. Linkers were further classified as independent domain linkers (IDL) when they connected two independent (non-interacting) domains and dependent domain linkers (DDL) when they connected two domains that interact with each other. Disordered regions overlapping with a single SCOP2 domain were classified as intra-domain loop (L), and disordered regions located at the start or end of a PDB chain were classified as termini (T). All regions not corresponding to IDLs, DDLs, L and T were classified as domains (D). Therefore, our definition of domains did not overlap strictly with the SCOP2 domain annotation since it was shaped by the identification of IDRs within PDB chains.

Before extracting IDRs, we noted that short, structured regions, such as isolated beta bridges (B) or very short helices (H) were embedded within IDRs and fragmented long IDRs into several shorter ones in our detection algorithm. To improve the representation of IDRs within our dataset, we performed several smoothing steps that converted short, isolated beta-bridges (B) and helices (H) into coils. However, smoothing within domains could result in the inclusion of residues that belong to short, structured regions into the intra-domain loops (L) dataset. To avoid this, we performed a two-step smoothing process that applies separate smoothing criteria for assignment of intra-domain loops (L) and domain linkers (IDLs/DDLs). In the first smoothing step (Smooth 1), we smoothed out all isolated beta-bridges (B) and helices (H) shorter than three residues, converting them into coils (C). Following this step, we extracted at least four continuous C residues as IDRs generating longer IDRs, akin to smoothing out short, structured regions. The IDRs generated from this step are designated as IDRs S1 and they will be used specifically for defining intra-domain loops (L), rather than domain linkers (IDLs/DDLs). We performed a second smoothing step (Smooth 2) on IDRs S1, smoothing out structured regions shorter than four residues without losing residue information on IDRs S1. Then again, we extracted at least four continuous C states as IDRs. The resulting IDRs were designated as IDRs S2. This second-step further extended the lengths of IDRs but was only applied for the extraction of domain linkers (IDLs/DDLs). Importantly, in both steps the smoothing preserved key features such as beta-strands involved in extensive hydrogen bonding with other structured regions within domains (labelled as E by DSSP) ensuring structural validity. The rationale for applying a second smoothing step to extract domain linkers but not to intra-domain loops is twofold: (a) allows capturing more and longer domain linkers, which are typically longer than intra-domain loops, and (b) prevents smoothing out structured regions within folded domains.

### Classification of loops and linkers

Following smoothing, we extracted domain linkers using IDRs S2 as the IDR annotation together with the SCOP2 domain boundaries (Supplementary File S2). First, we identified IDRs S2 that overlap with two adjacent SCOP2 domains and classified these as domain linkers. Second, we excluded IDRs from IDRs S1 that overlapped with these domain linkers. The remaining IDRs S1 overlapping with a single SCOP2 domain were classified as intra-domain loops (L) and those at the C or N-terminal regional as termini (T). Third, we refined SCOP2 domain annotations by excluding domain linkers and termini, resulting in a clear delineation of domain labels (D). Next, we revised the length of the domain regions and filtered out three PDB chains with domain regions shorter than 10 residues long that after manual inspection we noticed those correspond to small helices within a large IDR. Finally, domain linkers were parsed into independent (IDL) and dependent (DDL) domain linkers after classification of the domains by the following procedure:

For each case where an IDR connected two adjacent domains A and B, we calculated the number of contacts and the number of hydrogen bonds between the domains. For this, we measured the distance between the alpha-carbon (Cα) atoms of all possible amino acid pairs. A contact was counted when this distance was below the threshold (5 Å). The number of hydrogen bonds between domains was counted using DSSP annotations by using an energy threshold for hydrogen bond formation of −1 kcal/mol, which is higher than the default value of −0.5 kcal/mol established by DSSP [26]. Dependent domains were classified as domains containing contacts between three or more residue pairs and/or as domains containing two or more hydrogen bonds. Otherwise, domains were considered as independent.

Building upon this methodology, we have compiled the DLD dataset, which includes 1640 IDLs, 647 DDLs, 44896 L, 6044 D, and 2991 T distributed across 2646 PDB chains (Table 1). The DLD dataset is available at https://dld.chemeslab.org/, including PDB chains with residue-level region type annotations, as well as region type annotated with detailed information such as region start and end indices, length, amino acid sequence, PDB ID, PDB residue indexing, corresponding UniProt accession, and UniProt residue indexing.

### DisProt Dataset Collection

To analyze the quality of our dataset, we selected the DisProt Database. We downloaded from DisProt (version 9.5, December 2023) the Disordered Flexible Linkers (IDPO:00502) dataset, which contains 338 linkers across 267 protein sequences [10], offering a valuable benchmark for evaluating the quality and characteristics of our newly constructed DLD dataset. The term DisProt Linker is used in this work to refer to evidence of a disordered flexible linker region in DisProt.

### Sequence clustering analysis

We used MMSeq2 for clustering analysis. For the protein dataset, we include all unique sequences from the DLD and DisProt datasets or proteins containing at least one IDL, DDL or DisProt evidence longer than nine residues. For the region clustering analysis, we restricted the analysis to IDL, DDL or DisProt regions. Clustering was performed in the MMSeq2 [49]. Protein clustering was performed using the default (normal) clustering mode at 20% identity and 30% or 80% coverage, defining the minimal portion of sequence length overlap present in the query and target sequence (cov-mode: 0). Since regions are very short fragments, we modified the parameters for *k*-mers and coverage alignment to guarantee that short sequences within long sequences cluster together. Region clustering was performed with the following parameters for k-mers: k:5, spaced-kmer-mode:0, min-ungapped-score:10, mask:0, comp-bias-corr:0 and with the following parameters for alignment coverage: cov-mode:1, cluster-mode:2, single-step-clustering:1. Finally, region clustering was performed at 20% identity and 30% or 80% coverage.

### Sequence Embedding and Sequence Parsing

We used ProtTrans to perform sequence embeddings for data visualisation and classification of regions from the DLD dataset and DisProt Linkers from DisProt. To ensure that the embeddings incorporate broader sequence context and provide a more realistic representation of structural and functional environments, we first embedded the entire protein sequences rather than embedding only the isolated regions of interest. We then extracted the region embeddings from the sequence embeddings. Specifically, for a region such as an IDL containing *n* amino acids within a protein of *N* amino acids, we embedded the full sequence using ProtTrans, resulting in an *N* ∗ 1024 feature matrix. The embeddings corresponding to the n amino acids of the IDL region were then extracted (yielding an *n* ∗ 1024 matrix).

Following this strategy, we embedded 2646 PDB protein chains and generated the corresponding embedding matrices. From these matrices, we extracted the regions of interest, including IDLs, DDLs, Ls, Ds, and Ts from the DLD dataset. For comparison, we also embedded 267 DisProt sequences containing annotated linker regions and extracted their embeddings using the same methodology.

### Data Visualization using t-SNE plots

t-SNE in Tensorboard v2.17.1 is used to visualize the similarities and dissimilarities between different regions, including DLD regions and DisProt Linkers. To prepare the data for t-SNE, we first embed the protein sequences using the ProtTrans (as mentioned in the last section), which generates a 1024-dimensional vector for each residue. Since each region of interest (DisProt Linker, IDL, DDL, etc.) within the protein has its own length, the resulting embedded matrix has dimensions equal to the *lengt*ℎ *of t*ℎ*e region* ∗ 1024. To apply t-SNE, we compress each region into a single vector by calculating the mean of the embeddings across all residues within the region. This yields a 1 ∗ 1024 vector for each region, which is then passed to t-SNE for further dimensionality reduction to 2-dimensional (2D) space.

### Data Classification using M2O

To compare the similarities and dissimilarities between regions through binary classification, we categorized regions into two classes, class 0 and class 1. Then, the classifier, given a region as input, is trained to output a number between 0 and 1, indicating the likelihood of the region belonging to either class. For each pairwise classification task, we need a separate classifier. We evaluated two groups: In Group 1, we included four predictors comparing IDL, which is assigned to class 1 versus DDL, L, D or T regions, which are assigned to class 0, in each different classifier. Similarly, in Group 2 we included five predictors comparing in this case the DisProt linker dataset, which is assigned to class 1 against the IDL, DDL, L, D and T regions assigned to class 0 one in each predictor.

The classifiers are designed as a simple two-layer neural network, named M2O which means many-to-one (Supplementary Figure S13). M2O includes only the CNN layer as the learning component and a second layer for average pooling. The CNN layer, with a kernel size of 1, generates residue-level predictions. The number of input channels corresponds to the number of features, and the output channel is set to 1, representing the residue-level class prediction. Thus, the output of this layer is a vector with a length equal to the input region length. It is important to remark that the model does not consider the context of surrounding residues, as predictions are made independently for each position. The second layer averages the output from the first layer, producing a single value representing the entire region. This simple network architecture contains only 1025 parameters, all located in the CNN layer.

To assess the classifier performance reliably, we used the Area Under the Curve (AUC) metric, which is well-suited for imbalanced datasets. To further mitigate class imbalance, we adjusted the training sets to ensure similar class sizes (Supplementary Table S2, Training Size). We also applied 5-fold cross-validation to reduce variance due to any single train-test split. For example, in the task of distinguishing IDL from DDL regions, we clustered sequences using MMseqs2 [49] at 30% sequence identity and divided the resulting clusters into five groups. Each fold used four groups for training and one for testing, ensuring all data was evaluated once. We averaged the five resulting AUC scores to report the final model performance.

### Analysis of linker sequence and structure descriptors

Region sequence and structural analysis was performed using the local CIDER software package (version 0.1.21) [50] and SPARROW (version 0.2.2, under development) [51]. The sequence features calculated with CIDER includes: Fraction of positive, negative, polar, proline, aliphatic, aromatic, expansion and disorder promoting residues, mean Kyle-Dolittle hydrophobicity scaled to 0 and 9, and omega which is the patterning between charged/proline residues. The sequence features calculated with SPARROW include: The values of sequence charge decoration (SCD) which captures linear distribution of charged residues, sequence hydropathy decoration (SHD) which captures linear distribution of hydrophobicity and Kappa which captures the patterning between positive and negative charged residues. Predicted structural features calculated with SPARROW include: the end-to-end distance, radius of gyration, asphericity, and polymer law prefactor (ρ0) and scaling exponent (ν). Pfam domain annotations for each protein were run locally (Pfam release: 38.1). The AlphaFold models for each protein were downloaded from the AlphaFold database [52] in May, 2026. Additional protein annotations were collected from the Uniprot database and the full taxonomy was retrieved from the NCBI taxonomy database.

### DLD-Dataset web server development

The DLD-Dataset web server was developed in R using the Shiny framework version 1.13.0 [53] providing an interactive graphical interface for browsing and analyzing the IDLs, DDLs, termini and DisProt linkers. The web includes three-dimensional visualization implemented using the NGLVieweR package [54]. When available, the PDB chains and AlphaFold structural models are displayed together. PDB chains were previously structurally aligned to the corresponding AlphaFold model using BioPython version 1.87 [55] and Bio.PDB module [56] enabling direct comparison between experimental structures and predicted AlphaFold models within the web viewer. Interactive data visualizations were generated using the plotly package [57].

## AUTHOR ROLES

DM: Investigation, Formal analysis, Data curation, Writing – original draft; JG: Formal analysis, Data curation, Investigation, Visualization, Writing – original draft, Writing - review & editing, Resources, Software, Supervision of DM; HMGA: Formal analysis, Investigation, Visualization, Writing - review & editing. COL: Writing - review & editing, Resources, Software. GP: Supervision, Funding acquisition; LBC: Conceptualization, Supervision, Funding acquisition, Project administration, Writing – original draft. All authors reviewed and edited the final manuscript.

## DATA AVAILABILITY

Code is available at GitHub: https://github.com/deemeng/DLD-dataset/. The full DLD dataset is available at https://dld.chemeslab.org/.

