## Supplementary material for "Expanding the Landscape of Disordered Flexible Linkers: A Structural and Computational Framework for DLD dataset assembly": SI Figures and Tables

**This PDF file includes:**

Legends for Supplementary Files 1 and 2

Figures S1 to S13

Tables S1 and S2

Supplementary Text

**Supplementary File S1 (Separate File). SCOP domain definitions and classification.** This file was downloaded from the SCOP2 website (release 29-06-2022) and contains domain definitions along with the corresponding PDB and UniProt entry information. The fields are as follow for the SCOP family-level domain identifier (FA) and SCOP superfamily-level domain identifier (SF): SCOP domain identifier (DOMID), representative PDB ID for the family/superfamily domain (PDBID), domain region in PDB residue numbering (PDBREG), UniProt accession number (UNIID), domain region in the UniProt sequence (UNIREG). Last, the SCOP domain classification (SCOPCLA). Abbreviations include: TP = protein type, CL = protein class, CF = fold, SF = superfamily, FA = family.

**Supplementary File S2 (Separate File). SCOP Domain Boundaries in Multi-Domain PDB chains.** This file was derived by parsing Supplementary File S1 and contains detailed information on all SCOP domains identified within the 2867 PDB chains. For each PDB chain, additional metadata extracted from the PDB files is provided. The fields are as follow SCOP family-level domain identifier (FA-DOMID), representative PDB ID for the family domain (FA-PDBID), representative PDB chain ID (FA-CHAINID), family domain region in PDB residue numbering start index inclusive (FA-PDBREG-START) and end index inclusive (FA-PDBREG-END), UniProt accession number (FA-UNIID), family domain region in the UniProt sequence start index inclusive (FA-UNIREG-START) and end index inclusive (FA-UNIREG-END), unique identifier for each protein chain build as PDBI_CHAINID (seq_id), domain sequence length (length), experimental methodology used to determine the protein structure (experiment_type), and last, experimental resolution value reported in the PDB entry (resolution). Abbreviations include: XRay = X-ray crystallography, EM = electron microscopy, EC = electron crystallography, NMR = nuclear magnetic resonance spectroscopy, FA = family.

### Supplementary Figures

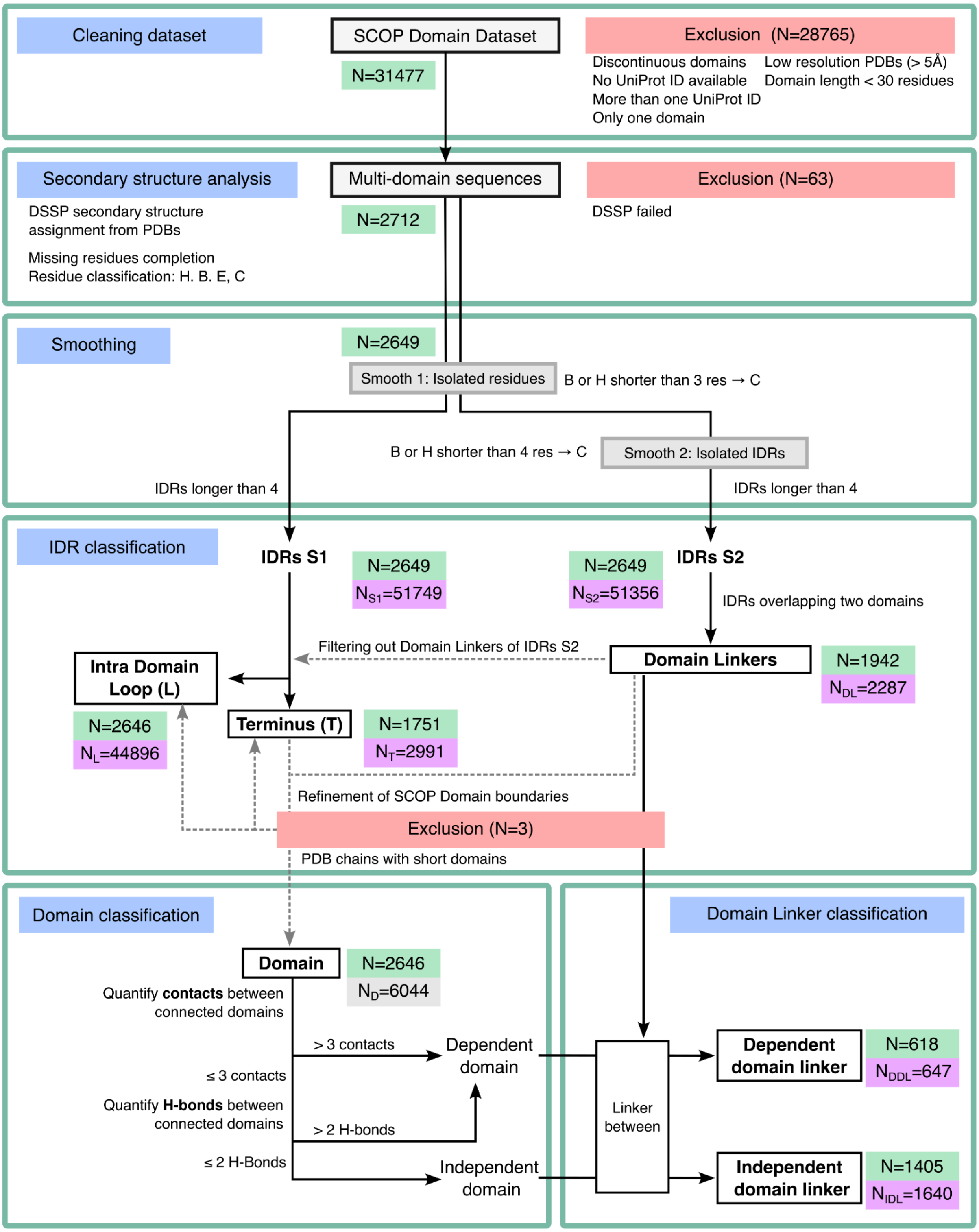


**Figure S1. Extended flowchart illustrating the construction process of the DLD dataset.** The numbers in the red Exclusion boxes are the number of PDB chains excluded. The numbers in green boxes indicate the number of PDB chains where IDRs or Domains are included. The number in purple boxes is the number of IDRs identified in each step. The number in gray boxes is the number of domain regions.

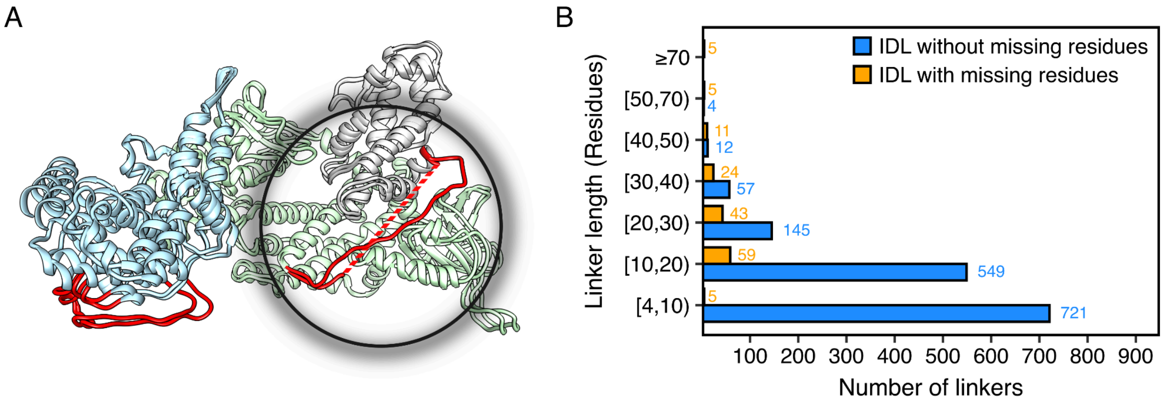


**Figure S2. Improvement of Independent Domain Linker (IDL) annotation with missing residue completion.** (A) Representative depiction of missing residue completion for the three-dimensional structure of the Son of Sevenless 1 protein (PDB 3KSY_A). Completing missing residues increases the length of the IDL in 17 residues. The effect of completing missing residues is shown by a structural alignment between 3KSY_A and the AlphaFold2 model (AF-Q07889-F1-model_v4) corresponding to this construct. For both representations, N and C- terminal regions are not displayed. Domains are colored in grey, green and blue from the N-terminus to the C-terminus. IDLs are highlighted in red. Dashed lines indicate missing residues. (B) Length distribution histogram for IDLs with or without missing residue completion. The total number of linkers for each category is indicated at the end of each bar. For linkers longer than 20 residues, completion of missing residues increased the total number by at least one-third. IDL: Independent domain linker.

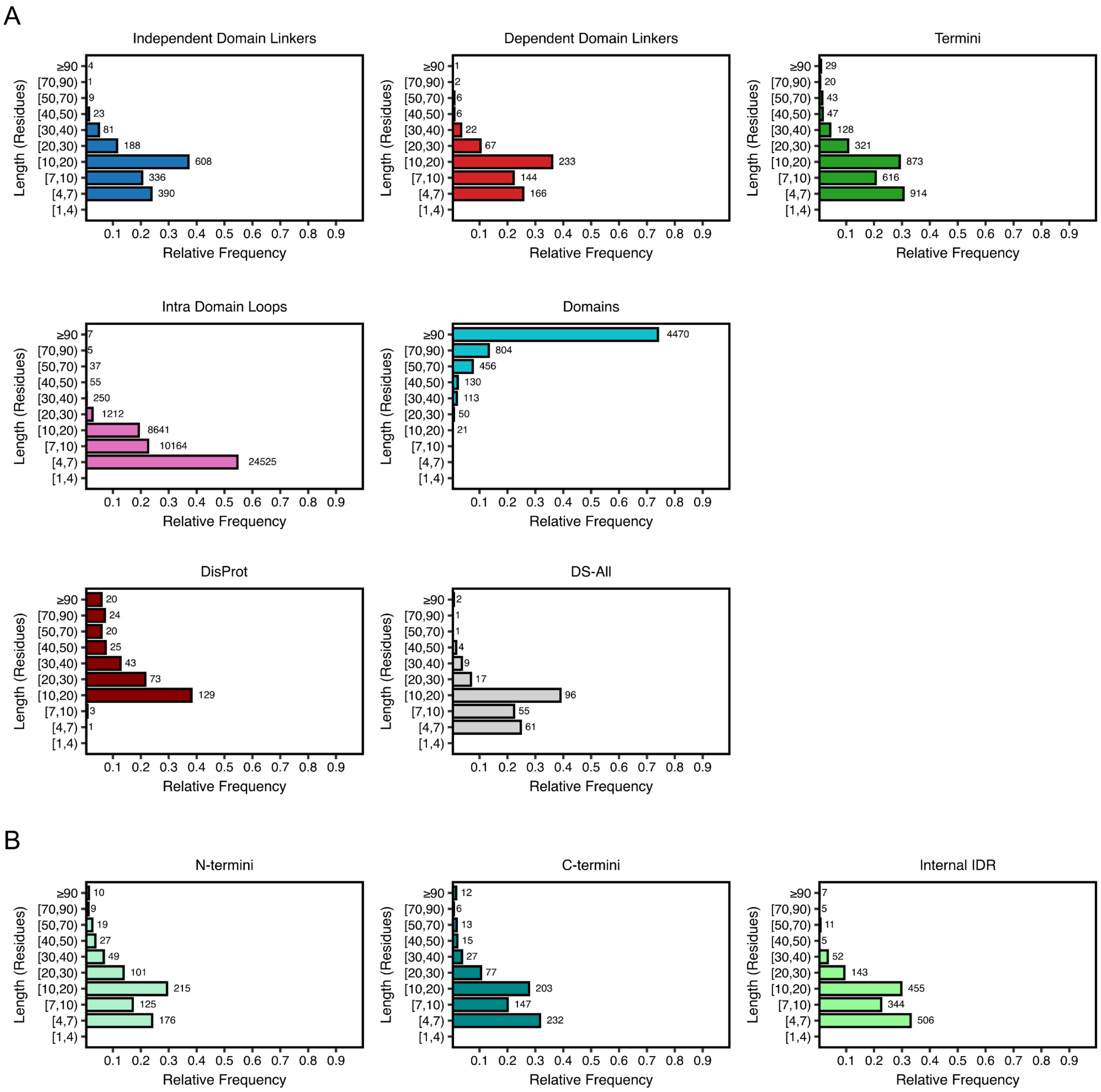


**Figure S3. Region and linker dataset length distribution.** (A) Linker length distribution for all datasets and region classes. (B) The termini region class was divided in the three groups according to region location in the full protein. The color of the bar indicates the region/linker dataset. The height of the bar indicates the relative frequency of the region/linker length. Lengths are grouped in intervals with different bin sizes. The number of regions/linkers in each interval is indicated at the top of the bar.

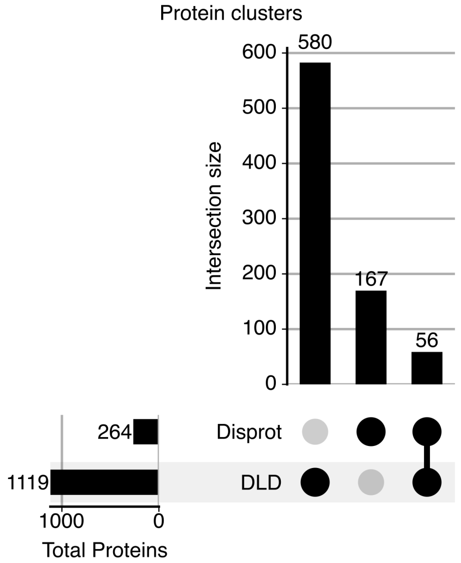


**Figure S4. Quantitation of overlap between the DLD and DisProt datasets.** Upset plot showing the number of DLD-specific, DisProt-specific and overlapping protein clusters. The datasets were filtered for regions > 9 residues and the thresholds used were 20% identity and 30% coverage. The side bar graph indicates the number of proteins for each dataset.

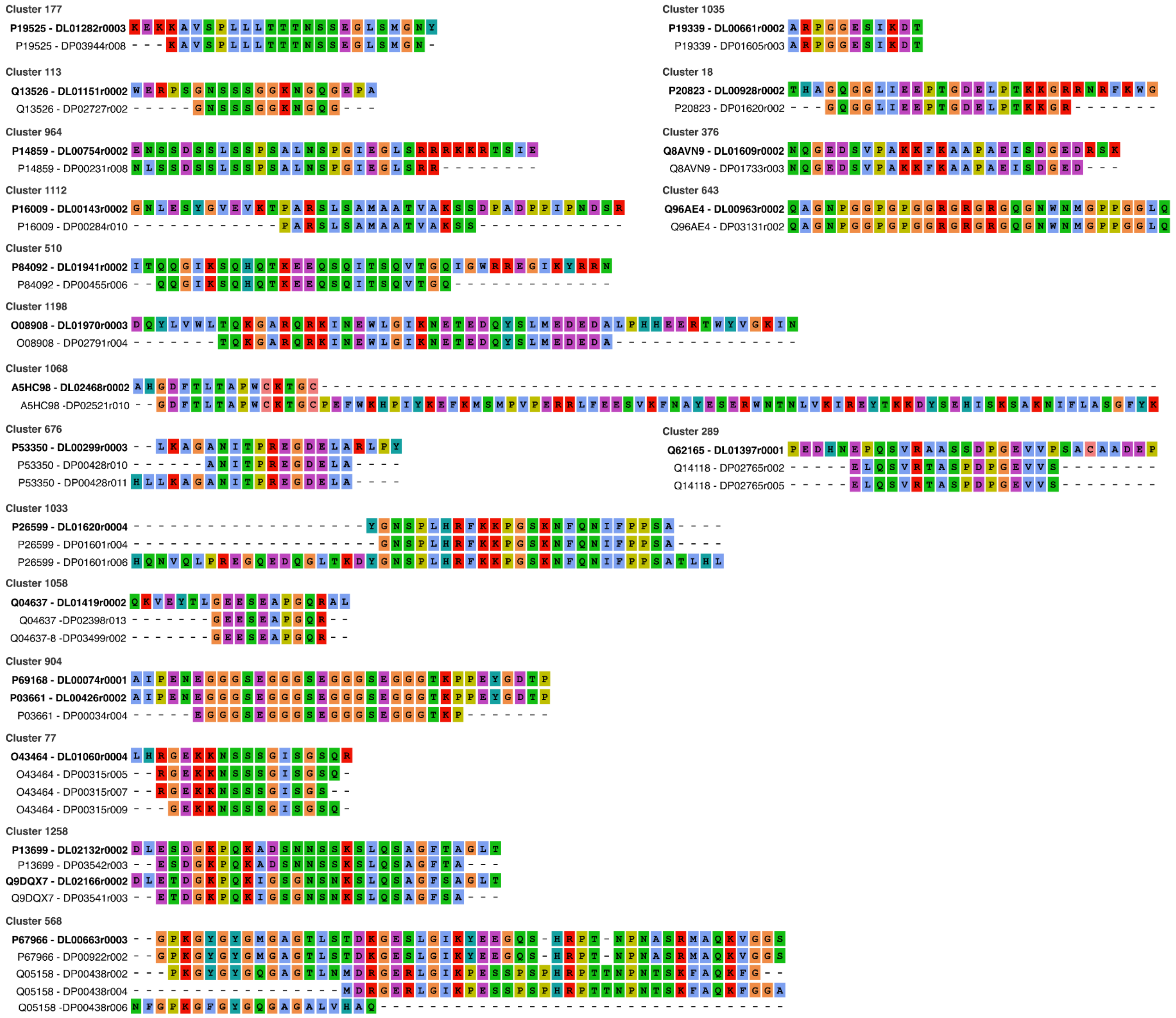


**Figure S5. Multiple sequence alignment of the 19 clusters shared between the DLD dataset regions and DisProt.** Highlighted in bold is the ID for the region in the DLD dataset. The clusters are sorted according to the size of the cluster. Sequence coloring follows the Clustal coloring code.

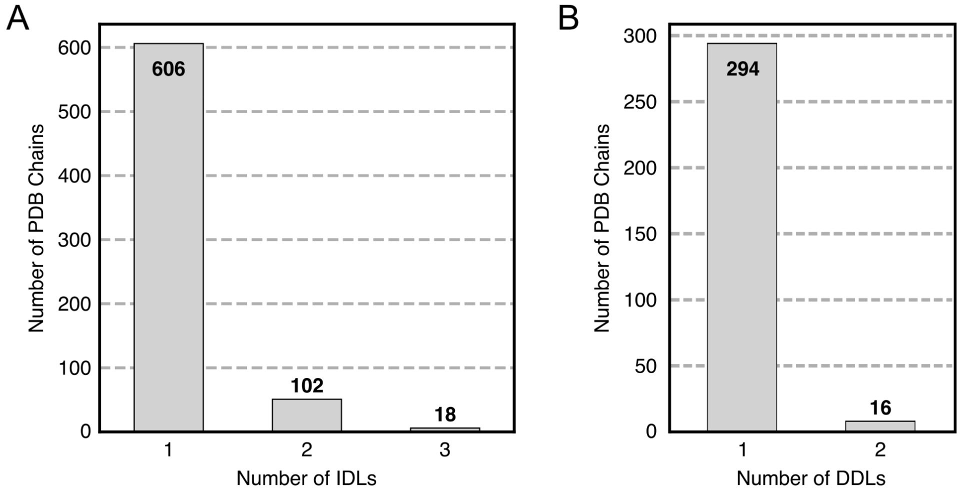


**Figure S6. Distribution of the number of short linkers per PDB chain.** A) Number of IDLs per PDB Chain for IDLs < 9 residues in length ($N_{TOTAL}$ = 726). B) Number of DDLs per PDB chain for IDLs < 9 residues in length ($N_{TOTAL}$ = 310). The number at the top of each bar indicates the total number of regions.

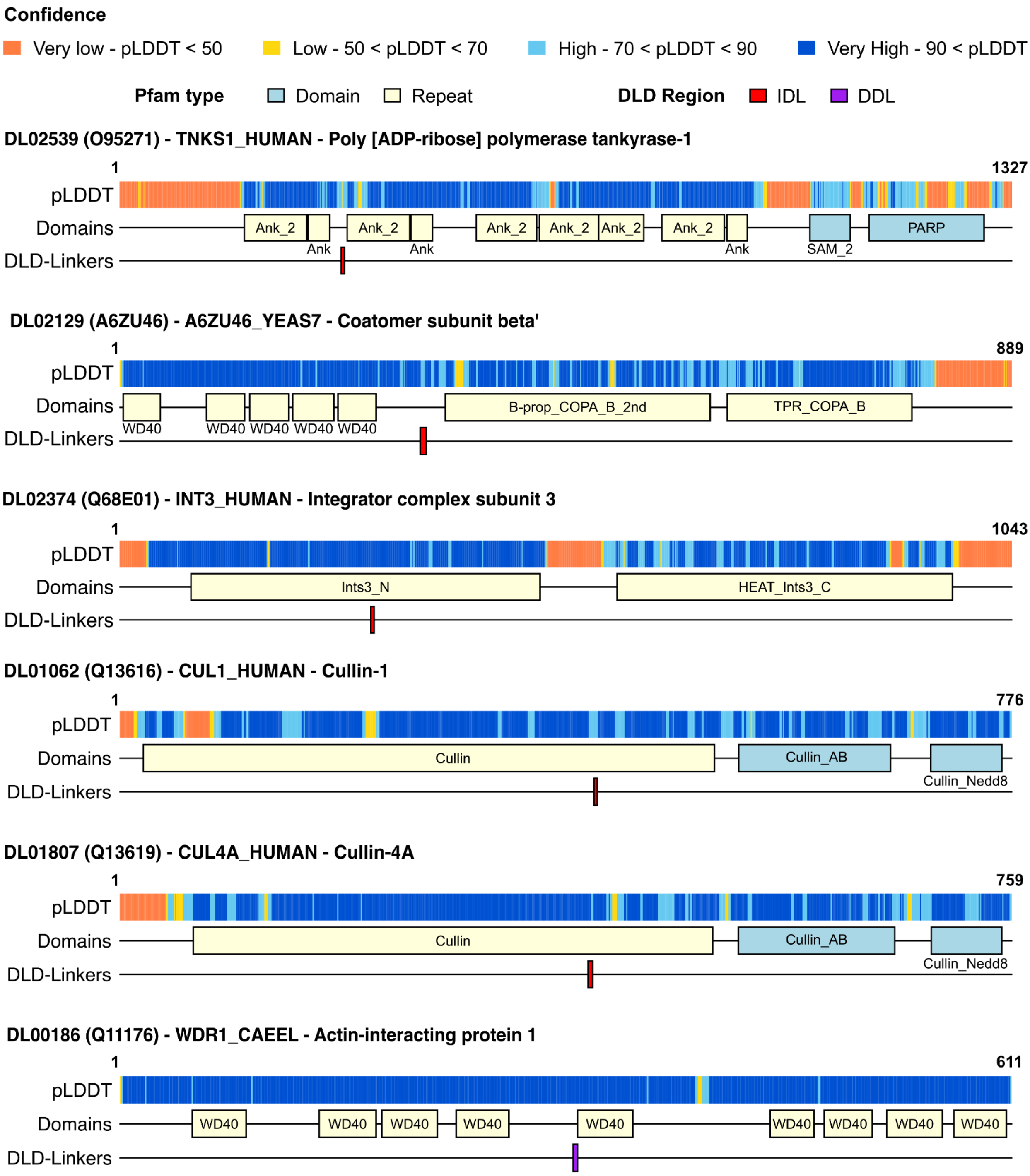


**Figure S7. Schematic representation of proteins with short linkers falling between or within repeat domains.** From top to bottom, the schematics show the AlphaFold2 pLDDT prediction, Pfam domain annotations and DLD-Linkers (four IDLs and one DDL). Pfam domain annotations are coloured based on the Pfam type (pale yellow: repeat, light blue:domain).

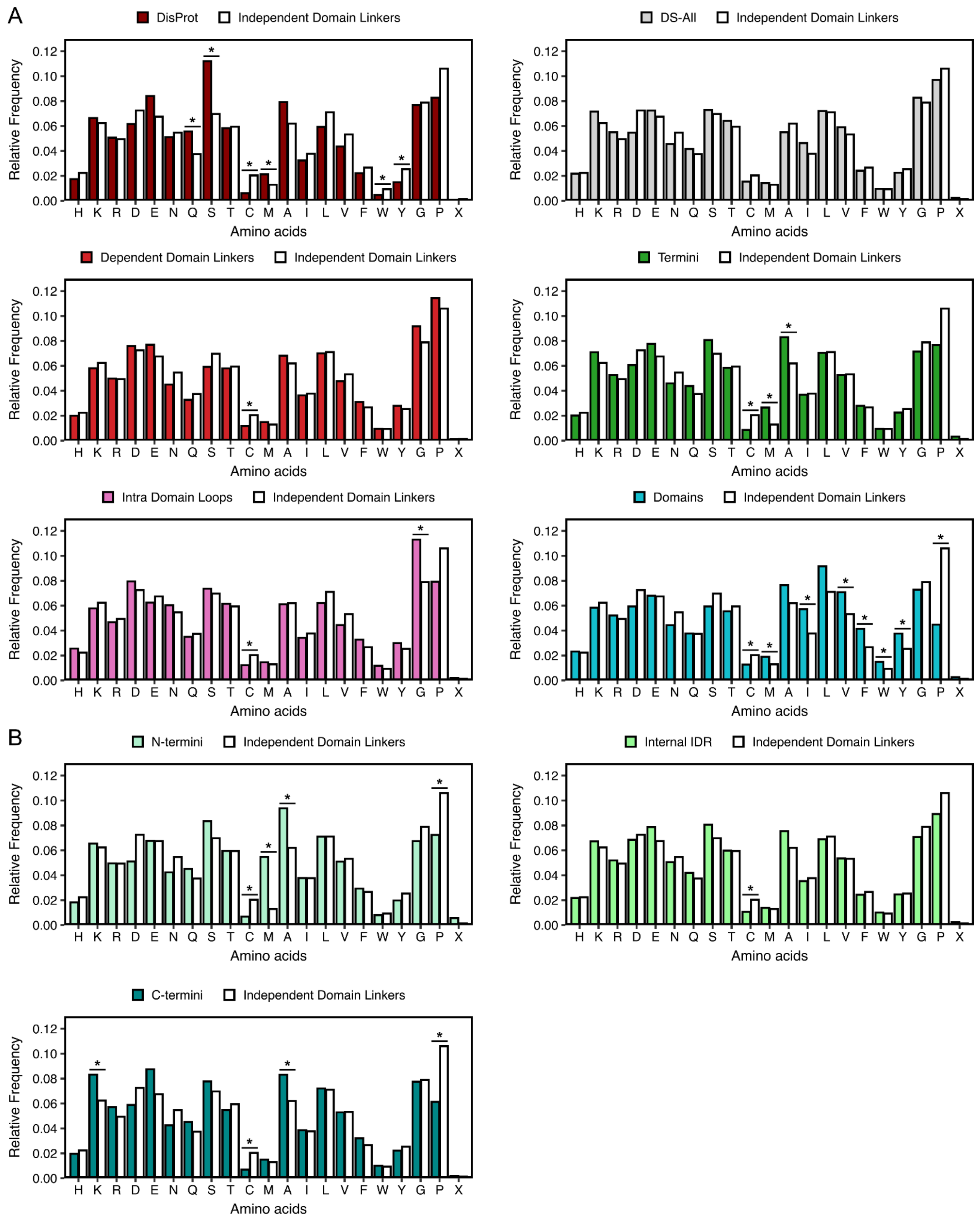


**Figure S8. Sequence composition histograms for regions and linkers.** (A) Sequence composition comparison between all regions/linker dataset and the independent domain linkers. (B) Sequence composition comparison between the N-termini, Internal IDRs and C-termini subsets from the termini region class and the independent domain linkers. The height of the bar indicates the relative frequency of the amino acid in the region/linker dataset. For the DS-All dataset, we only determined sequence composition for PDBs where the chain id was annotated. * Indicates that the difference between the relative frequency of each amino acid in the selected class is at least 30% lower or higher compared to the relative frequency found in the IDLs.

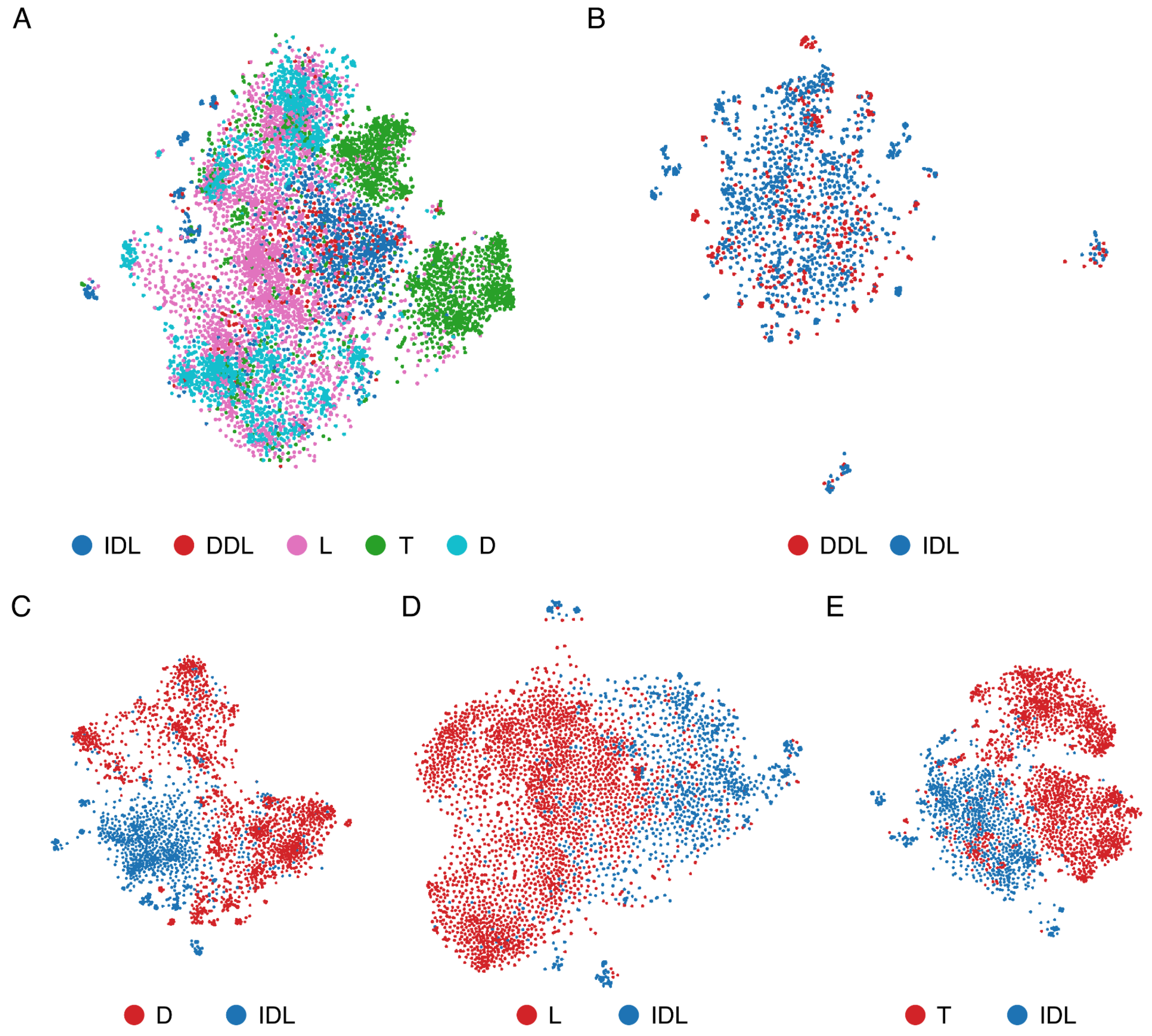


**Figure S9. t-SNE plots for pair comparisons in the DLD dataset.** (A) t-SNE plot of the 5 regions in the DLD dataset. (B-E) Pairwise comparison t-SNE plots. (B) DDL vs IDL, (C) D vs IDL, (D) L vs IDL and (E) T vs IDL. DLD: Domain-Linker-Domain. IDL: Independent domain linker. DDL: Dependent domain linker. L: Intra domain loop. T: Terminus. D: Domain.

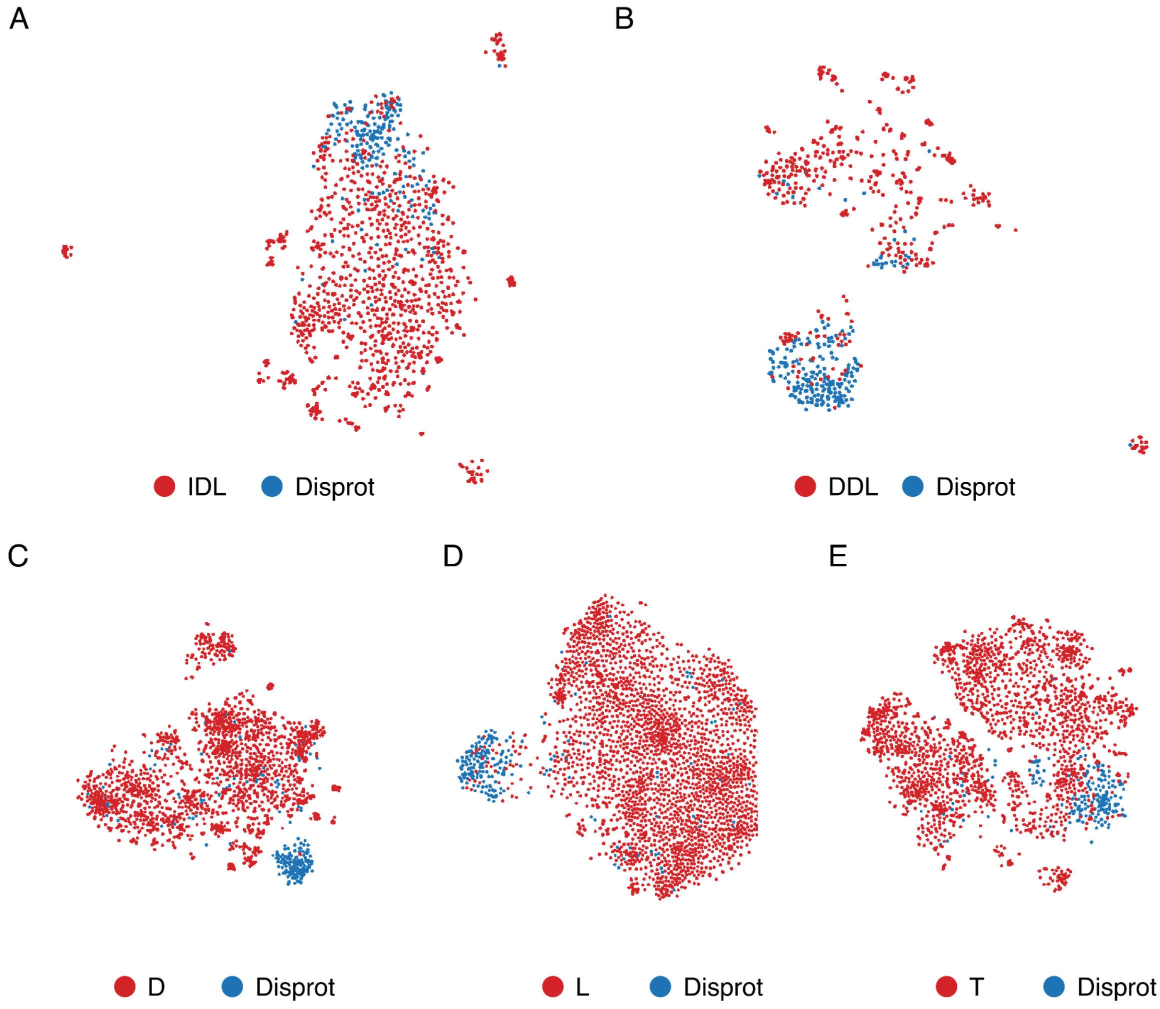


**Figure S10. t-SNE plots for pair comparisons between regions in the DLD dataset and the DisProt linker dataset.** (A) IDL, (B) DDL, (C) D, (D) L and (E) T regions vs DisProt linkers. DLD: Domain Linker Domain. IDL: Independent domain linker. DDL: Dependent domain linker. L: Intra domain loop. T: Terminus. D: Domain.

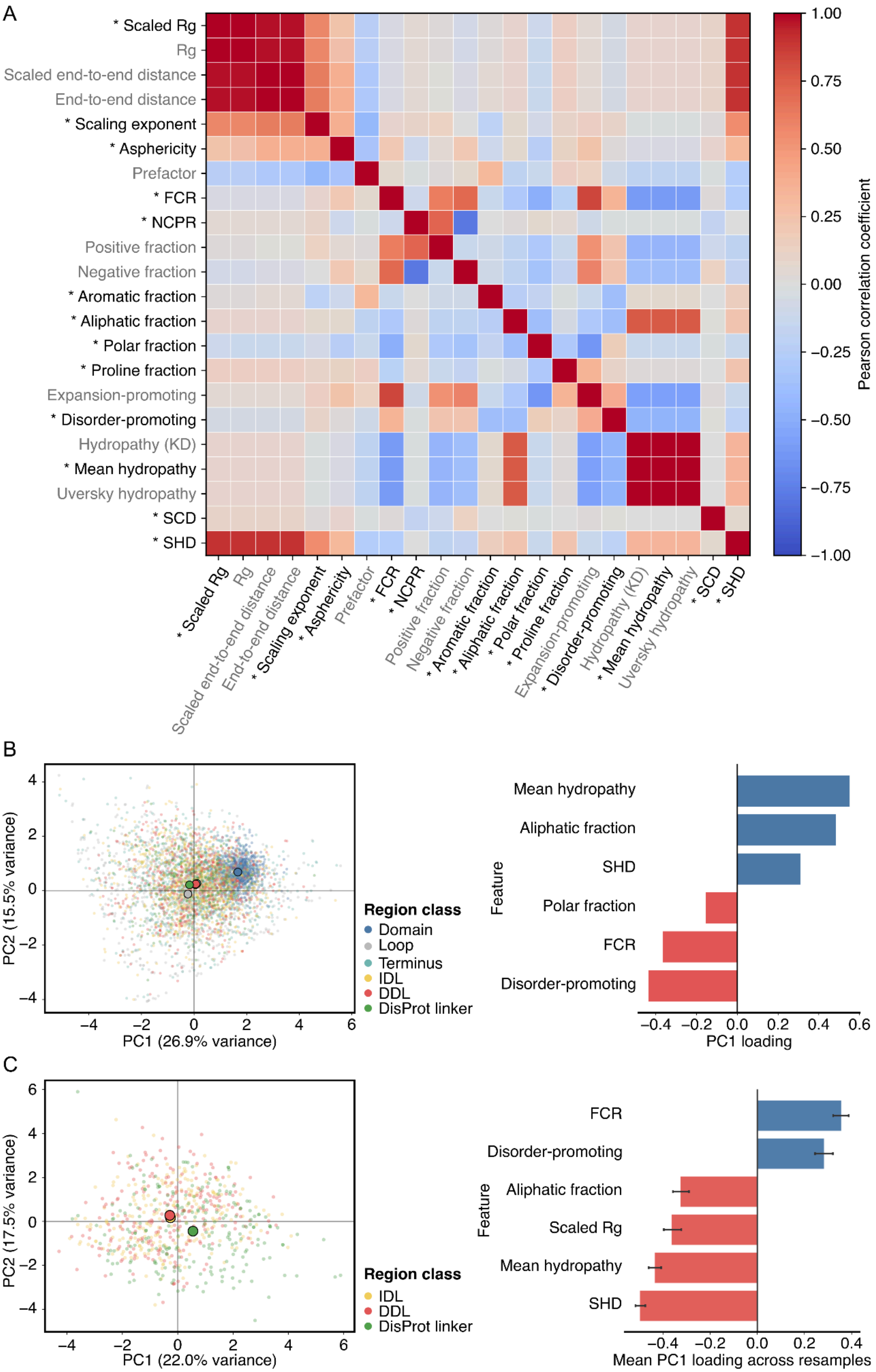
A
* Scaled Rg
Rg
Scaled end-to-end distance
End-to-end distance
* Scaling exponent
* Asphericity
Prefactor
* FCR
* NCPR
Positive fraction
Negative fraction
* Aromatic fraction
* Aliphatic fraction
* Polar fraction
* Proline fraction
Expansion-promoting
* Disorder-promoting
Hydropathy (KD)
* Mean hydropathy
Uversky hydropathy
* SCD
* SHD
1.00
0.75
0.50
0.25
0.00
−0.25
−0.50
−0.75
−1.00
Pearson correlation coefficient
B
PC2 (15.5% variance)
PC1 (26.9% variance)
Region class
Domain
Loop
Terminus
IDL
DDL
DisProt linker
Mean hydropathy
Aliphatic fraction
SHD
Polar fraction
FCR
Disorder-promoting
Feature
PC1 loading
C
PC2 (17.5% variance)
PC1 (22.0% variance)
Region class
IDL
DDL
DisProt linker
FCR
Disorder-promoting
Aliphatic fraction
Scaled Rg
Mean hydropathy
SHD
Feature
Mean PC1 loading across resamples

**Figure S11. Principal component analysis of sequence-structure derived features for the DLD and DisProt datasets.** (A) Correlations among candidate sequence and structural features. Pearson correlations were calculated across the common complete-case set of 50,500 non-domain regions, including loops, termini, IDLs, DDLs, and DisProt linkers. Asterisks indicate the 13 features retained for subsequent principal component analyses. (B) Sequence-composition PCA of all regions. *Left.* Principal component analysis of ten sequence-derived features across folded domains, loops, termini, IDLs, DDLs, and DisProt linkers. For visualization, up to 1,000 regions were sampled per class; large outlined markers indicate centroids calculated from all regions in each class. *Right.* PC1 loadings for the six features with the largest absolute contributions. (C) Sequence and structural-composition PCA of IDLs, DDLs and DisProt linkers. *Left.* Representative PCA of IDLs, DDLs, and DisProt linkers using 13 sequence and structure derived features. Points represent individual regions and large outlined markers indicate class centroids. *Right.* Mean PC1 loadings across the 500 matched resamples for the six features with the largest absolute mean loadings; error bars indicate empirical 95% intervals. PC1 was sign-aligned in each resample so that DisProt linkers had higher mean PC1 scores than the pooled IDL and DDL groups.

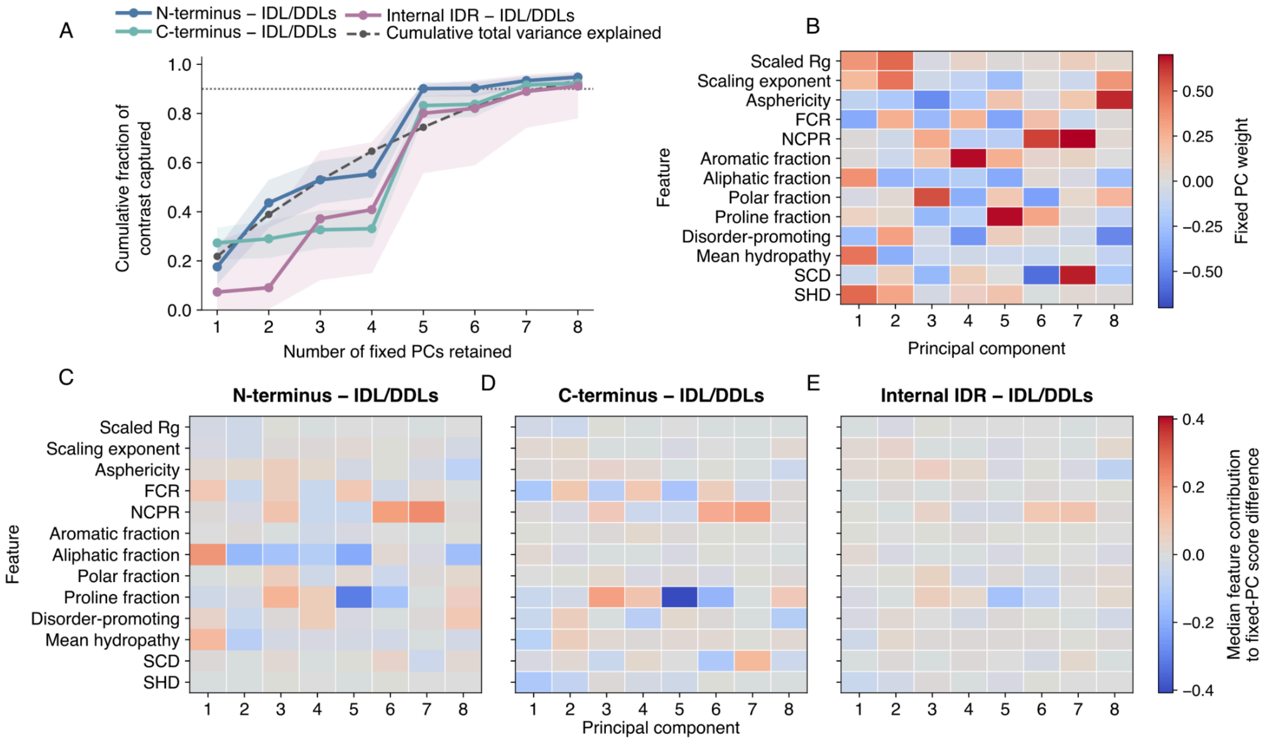


**Supplementary Figure S12. Fixed-PCA interpretation of feature differences between terminal regions, Internal IDRs, and structural linkers.** (A) Cumulative fraction of the squared standardized feature difference captured by the first eight fixed principal components for N-termini, C-termini, and Internal IDRs relative to the equally weighted IDL/DDLs reference group across 500 length-matched resamples. Solid lines show median cumulative capture and shaded regions show the empirical 2.5th–97.5th percentile range across resamples. The dashed line indicates cumulative total variance explained by the fixed PCA, and the horizontal dotted line marks 90% contrast capture. (B) Feature weights of the first eight components of the fixed PCA, fitted to the pooled matched regions across all resamples. (C–E) Median feature-wise terms contributing to the fixed-PC score difference between the indicated region class and the IDL/DDLs reference group. For each resample, the contribution of feature *j* to principal component *k* was calculated as the standardized feature difference for feature *j* multiplied by its fixed weight on component *k*. Heatmap values are the median of these terms across resamples. Positive and negative values indicate contributions in the positive and negative directions, respectively, of the corresponding fixed PC. All three heatmaps use the same feature order and symmetric color scale.

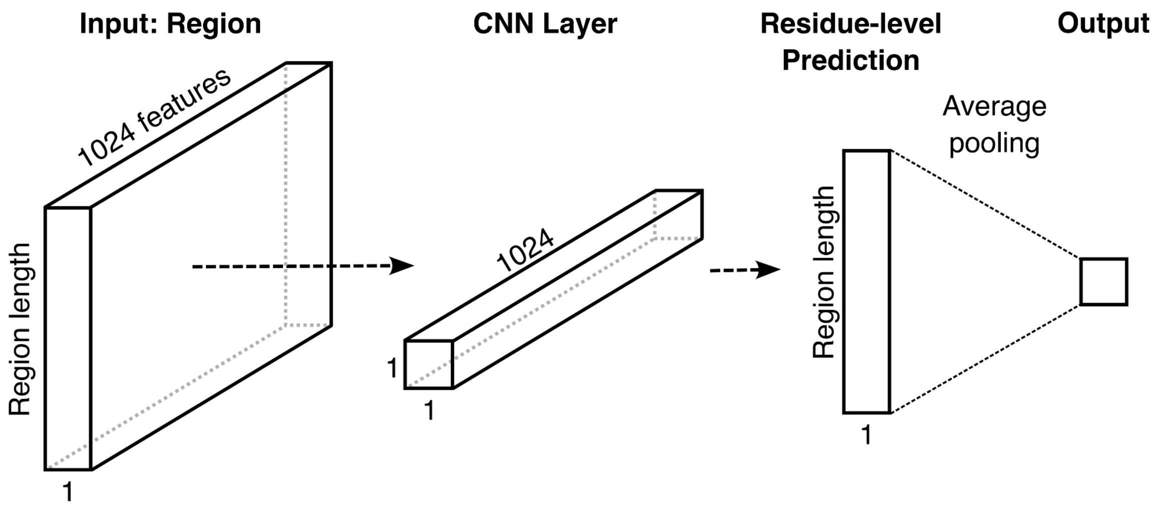


**Figure S13. Model architecture of the M2O classifier.** The only learning layer (with weights to learn through training) is the CNN layer. Residue-level prediction is the output of the CNN layer, then it passes to Average Pooling which calculates the mean of the Residue-level prediction.

**Supplementary Tables**

**Table S1. Clustering of the DLD and DisProt datasets at the protein and region level.**

| | | | Protein clustering at 20 % identity* | | Region clustering at 20 % identity* | |
|---|---|---|---|---|---|---|
| | **Unique Proteins** | **Unique Regions** | **80% Cov.*** | **30% Cov.*** | **80% Cov.*** | **30% Cov.*** |
| **All regions** | | | | | | |
| **# DLD†** | 1897 | 2287 | 1185 | 917 | 2075 | 2069 |
| **# DisProt** | 234 | 338 | 193 | 166 | 276 | 268 |
| **# DLD† & DisProt** | 33 | - | 48 | 61 | 18 | 19 |
| **Regions⨍ > 9 res.** | | | | | | |
| **# DLD†** | 1086 | 1251 | 727 | 580 | 1093 | 1087 |
| **# DisProt** | 231 | 334 | 195 | 167 | 272 | 264 |
| **# DLD & DisProt** | 33 | - | 43 | 56 | 18 | 19 |

*We used MMSeq2 for clustering with a minimum sequence identity of 20% and 80% or 30% sequence alignment coverage defining the minimal portion of sequence length overlap for grouping sequences together. For region clustering, the sequence alignment coverage was required for the longest sequence in the cluster.

† DLD denotes the sum of proteins/regions containing IDLs and proteins/regions containing DDLs.

⨍Protein clustering was performed on the DLD dataset restricted to proteins containing IDLs/DDLs longer than 9 residues and DisProt clustering was performed on proteins containing evidences longer than 9 residues.

**Table S2. Classification analysis for pair comparisons between linker regions and other regions.**

| Region 1 | Region 2 | Predictor | Training size Region 1, Region 2 | Training epoch | AUC |
|---|---|---|---|---|---|
| **IDL Group** | DDL | pIDL_DDL | 1640, 647 | 600 | **0.762** |
| | L | pIDL_L | 1640, 1640 | 150 | 0.966 |
| | D | pIDL_D | 1640, 1640 | 150 | 0.999 |
| | T | pIDL_T | 1640, 1640 | 150 | 0.974 |
| **DisProt Linker Group** | IDL | pIDPO_IDL | 338, 1640 | 150 | **0.875** |
| | DDL | pIDPO_DDL | 338, 647 | 150 | 0.937 |
| | L | pIDPO_L | 338, 1640 | 150 | 0.983 |
| | D | pIDPO_D | 338, 1640 | 150 | 0.992 |
| | T | pIDPO_T | 338, 1640 | 150 | 0.946 |

### Supplementary Text

#### Principal component analysis of sequence- and structure-derived features

To characterize interpretable properties associated with the different regions, we calculated sequence- and predicted structure-derived features for all regions in the DLD and DisProt datasets (Table 1).

Sequence-derived features were calculated using LocalCIDER and SPARROW. The sequence features calculated with CIDER include: Fraction of positive, negative, polar, proline, aliphatic, aromatic, expansion and disorder promoting residues, two different values of hydropathy (mean hydropathy and Uversky hydropathy) and omega which is the patterning between charged/proline residues. The third value of hydropathy (hydropathy KD), the values of sequence charge decoration (SCD) which captures linear distribution of charged residues, sequence hydropathy decoration (SHD) which captures linear distribution of hydrophobicity and Kappa which captures the patterning between positive and negative charged residues were calculated using SPARROW as well as the predicted structural features that include the two options, scaled or not of end-to-end distance and radius of gyration, and finally, asphericity, and polymer law prefactor ($\rho 0$) and scaling exponent ($v$). The values for Kappa and Omega were not possible to calculate for a high proportion of loops and they were discarded in the analysis. Since the predictor for structural features in SPARROW is trained for disordered regions, the structural features were not included in the analysis of folded domains. To reduce redundancy among features, pairwise Pearson correlations were calculated on the common complete-case set of 50,500 non-domain regions, comprising loops, termini, IDLs, DDLs, and DisProt linkers. Where highly correlated features represented the same feature group, a single representative variable was retained. In particular, mean Kyte–Doolittle hydropathy was retained among the hydropathy measures, and scaled radius of gyration was retained to represent the highly correlated polymer features. The resulting reduced feature set comprised 13 variables: fraction of charged residues (FCR), net charge per residue (NCPR), aromatic fraction, aliphatic fraction, polar fraction, proline fraction, disorder-promoting composition, mean hydropathy, SCD, SHD, scaled radius of gyration, scaling exponent, and asphericity. Region length was retained for matching procedures but was not included as an input variable in PCA.

Two PCA analyses were performed. First, sequence-composition differences across all region classes were assessed using the ten sequence-derived features available for both domains and non-domain regions. PCA was performed on folded domains, loops, termini, IDLs, DDLs, and DisProt linkers. Second, IDLs, DDLs, and DisProt linkers of at least 10 residues were compared using the full set of 13 sequence- and structure-derived features. Termini were excluded because a substantial proportion did not correspond to the true N- or C-terminus of the protein. The resulting dataset comprised 914 IDLs, 337 DDLs, and 334 DisProt linkers. Because DisProt linkers had a different length distribution from IDLs and DDLs, length was controlled through repeated matched resampling. Specifically, regions were assigned to seven bins based on log-transformed length, and 500 matched subsets were generated by sampling equal numbers of regions from each class within each bin. Each matched subset contained 225 IDLs, 225 DDLs, and 225 DisProt linkers. PCA was independently performed for each matched subset using the 13 features, without including region length as an input feature. For visualization, a representative resample was selected based on the centroid distance between DisProt linkers and the pooled IDL-DDL group being closest to the median centroid distance across the 500 resamples. PC1 loading signs were aligned across resamples such that DisProt linkers had higher mean PC1 scores than the pooled

IDL and DDL groups. Mean PC1 loadings and empirical 95% intervals were then calculated across the 500 length-matched resamples.

To further examine the heterogeneous termini class, termini were subdivided into bona fide N-termini and C-termini, and internally located regions referred to as Internal IDRs. These categories were compared with IDLs and DDLs using the same reduced set of 13 sequence- and structure-derived features. To control for length and class-size differences, 500 repeated matched resamples were generated by assigning regions to seven log-transformed length bins and sampling equal numbers of IDLs, DDLs, N-termini, C-termini, and Internal IDRs within each bin. Each matched subset contained 288 regions per class, 1,440 regions in total.

A single scaler and PCA were fitted to the pooled matched regions from all 500 resamples, providing a common fixed PCA coordinate system. For each resample, features were transformed using this fixed scaler, and all sampled regions were projected onto the fixed PCA axes. Centroids were then calculated for each category, and the structural-linker reference was defined as the equally weighted mean of the IDL and DDL centroids. Direct feature contrasts were calculated within each resample by subtracting this IDL/DDLs reference centroid from the N-terminus, C-terminus, or Internal IDR centroid, yielding feature differences in standardized units for each comparison. For each feature, the median difference and the empirical 2.5th–97.5th percentile range were then summarized across the 500 resamples.

For visualization of the PC1–PC2 projection, the representative resample was selected as the one whose ten pairwise centroid distances among the five classes were closest to the median centroid-distance profile across the 500 resamples. To assess how each multifeature contrast was represented in fixed PCA space, the contrast vector for each comparison was projected onto each fixed PC, yielding the fixed-PC score difference between the indicated category and the IDL/DDLs reference group along that component. These score differences were used to calculate cumulative contrast capture: the fraction of contrast captured by the first $k$ PCs was defined as the sum of squared fixed-PC score differences from PC1 to PC$k$, divided by the total squared length of the contrast vector in standardized feature space. To visualize which features contributed to the projected score differences, each fixed-PC score difference was decomposed into feature-wise terms by multiplying the standardized difference for each feature by the corresponding fixed PC weight. Median cumulative contrast capture and median feature-wise PC contributions were summarized across the 500 resamples.
